# DNA replication and polymer chain duplication reshape the genome in space and time

**DOI:** 10.1101/2024.03.12.584628

**Authors:** Dario D’Asaro, Maxime M. C. Tortora, Cédric Vaillant, Jean-Michel Arbona, Daniel Jost

## Abstract

In eukaryotes, DNA replication constitutes a complex process whereby multiple origins are stochastically fired, and from which the replication machinery proceeds along chromosomes to achieve the faithful synthesis of two identical copies of the genome during the S-phase of the cell cycle. Experimental evidence show a functional correlation between the dynamics of replication and the spatial organization of the genome inside cell nuclei, suggesting that the process of replicating DNA may impact chromosome folding. However, the theoretical and mechanistic bases of such an hypothesis remain elusive. To address that question, we propose a quantitative, minimal framework that integrates the dynamics of replication along a polymer chain by accounting explicitly for the progression of the replication machinery and the resulting formation of sister chromatids. By systematically characterizing the 3D structural consequences of replication, and of possible interactions between active replication machineries, we show that the formation of transient loops may potentially impact chromosome organization across multiple temporal and spatial scales, from the level of individual origins to that of the global polymer chain. Comparison with available microscopy and chromosome conformation capture data in yeast suggests that a replication-dependent loop extrusion process may be acting *in vivo*, and may shape chromosomes as loose polymer bottle-brushes during the S-phase. Lastly, we explore the post-replication relative organization of sister chromatids and demonstrate the emergence of catenations and intertwined structures, which are regulated by the density of fired origins.

## I. INTRODUCTION

Inside the eukaryotic nucleus, DNA is tightly packed into a polymer-like structure called chromatin. Understanding chromatin self-assembly and its higher-order 3D architecture [1–3] within the confined space of the nucleus constitutes a major challenge in biology as chromatin organization has key functional roles in many fundamental biological processes such as gene transcription, genome replication and DNA repair [3, 4]. In particular, the complex 3D genome organization must exhibit strong plasticity to assist the functional and structural changes observed during the cell cycle [5, 6]. A remarkable illustration of this is observed during the S phase, when chromosomes are progressively duplicated before cell division, while the 3D chromatin architecture must adapt to the presence of newly replicated sister chromatids. This process is finely regulated in order to achieve the maintenance of cell identity through the faithful inheritance of all genetic information.

Eukaryotic replication is initiated at multiple genomic loci [7], called origins of replication, that are licensed during the G1 phase and later activated throughout the S phase via the combined action of several replication-associated factors [8, 9]. Origins are fired with different efficiencies in a non-trivial timing, generally resulting in complex, heterogeneous temporal patterns of replication along the genome [10]. Several coupled analyses of the 3D genome in interphase and the timing of replication along the genome have demonstrated strong correlations between the two [11, 12]. For example, early replicating domains coincide with active — open — spatial compartments while late replicating domains with more condensed — repressed — compartments. However, while these two processes must be closely interconnected, little is known regarding their causal relationships. On the one hand, the observed correlation suggests that the pre-existing spatial organization may be involved in the regulation of firing events at origins. In this picture, local accessibility to chromatin may play a prominent role in governing the heterogeneous firing of the different origins. On the other hand, the replication dynamics, usually defined within the 1D context of the genomic sequence, occurs concurrently with possible changes in 3D genome organization. Indeed, several works using (super-resolution) microscopy and chromosome conformation capture (3C) techniques have shown a reorganization of chromosomes while the genome is being replicated during S phase [5, 13–17]. Early microscopy studies suggested that each pair of replication machineries originating from the same origin, forming a so-called replicon, and replicating DNA in opposite directions may co-localize in 3D space [17], and that, more generally replication machineries may segregate inside specific nuclear bodies called replication factories [14, 18, 19]. However, more recent studies were able to resolve single replicons within such condensates, suggesting that factories may actually naturally emerge from the pre-existing 3D chromosome organization [15, 16] and challenging the idea of functional self-assembly of replicons. In addition, 3C techniques applied over the whole cell-cycle, have observed some reorganization happening during the S-phase [5, 13]. However, it was unclear if such changes emanate from the replication process itself or from other mechanisms acting on chromatin, like cohesin- or condensin-mediated loop extrusion, that may also act during S-phase. Overall, the current body of experimental data is therefore insufficient to provide a clear mechanistic picture of how replication may impact genome organization.

Over the years, polymer physics has proven to be a powerful framework to get new insights into the physico-chemical mechanisms driving chromosome organization and dynamics, over a wide range of spatial and temporal scales [20]. However, to date, most theoretical approaches have focused on genome folding during the G1 phase[21, 22] and, to a lesser degree, on late G2 phase and mitosis [23–26]. Furthermore, although several modelling studies have previously tackled the 1D dynamics of eukaryotic replication [27–29], a theoretical investigation of its implications on 3D structure remains largely lacking. Recent polymer models have investigated the replication process in the context of circular and highly confined bacterial chromosomes [30, 31], highlighting the role of entropic repulsion triggered by the various loops formed during replication. However these studies mostly focus on the segregation of circular chromosomes, carrying a single origin — while replication in eukaryotic linear genomes requires multiple origin firings, and segregation mainly occurs when replication has been completed. Therefore, aiming to fill this gap, we develop a computational framework to describe the spatio-temporal dynamics of chromatin during S phase by accounting for the explicit duplication of a linear polymer chain. The active 3D process of DNA replication further offers the opportunity to theoretically investigate the out-of-equilibrium mechanics of polymer chain self-duplication starting from specific sites, and to better understand how the replication process may dynamically affect the local and global spatial properties of both the template linear polymer and its sister chromatid. Experimental and theoretical studies have shown chromosome 3D organization results in part from the activity of structural maintenance complexes such as cohesins and condensins actin a DNA loop extruders [5, 26, 32]. By analogy, we may expect the colocalisation of replication machineries in a single replicon to work as an extrusion like process (Fig. 1D) and to structure the sister chromatids. We aim with our model to challenge such a hypothesis and characterize systematically the role of replicon properties in genome organization.

**FIG. 1.**
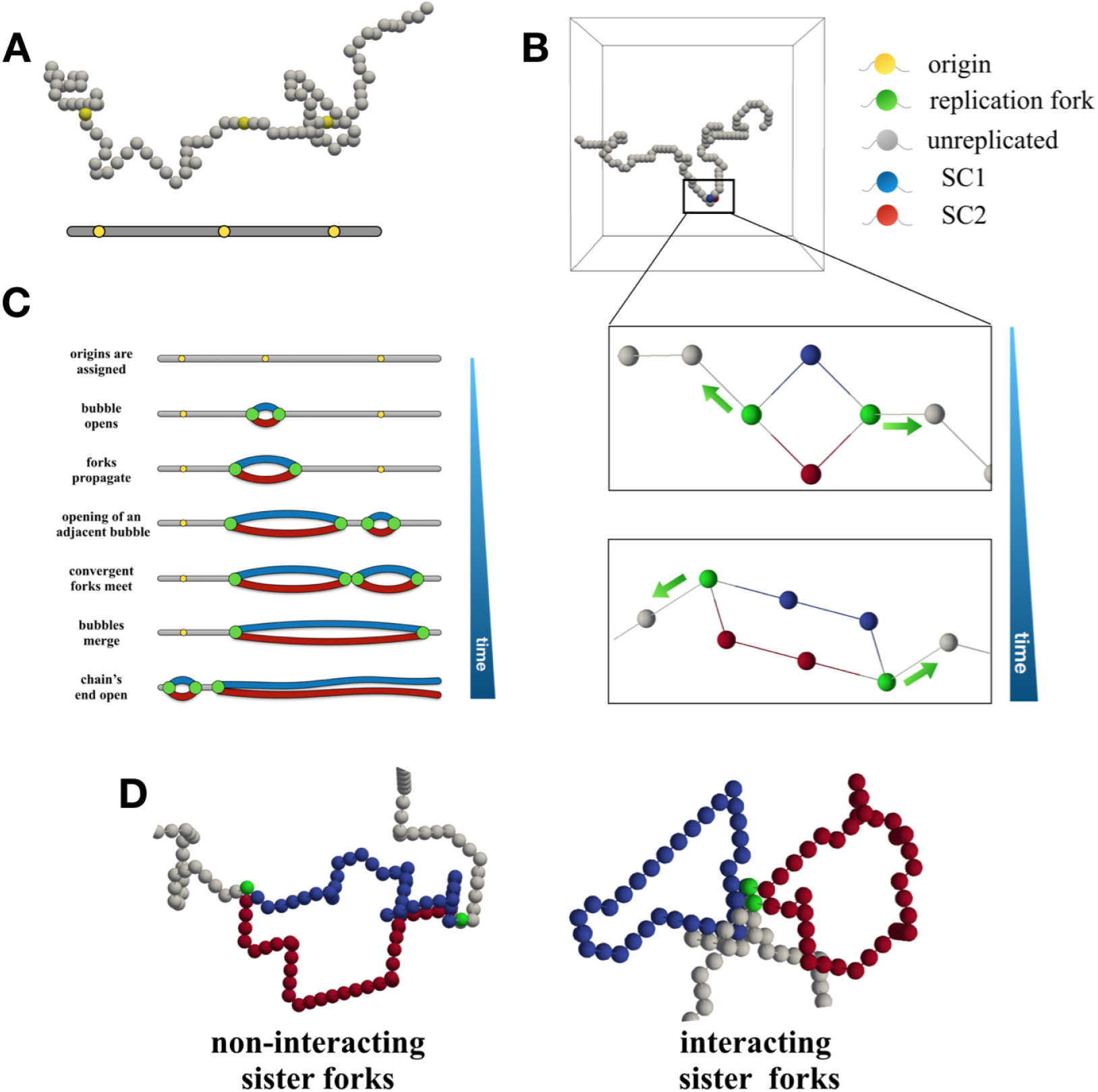
Model of self-replicating polymer. (A) Example of a template polymer chain where a set of monomers were selected as origins of replication (beads in yellow). (B) Example of a firing event and fork progression. After firing, a new monomer is inserted and connected to the chain creating a replication bubble leading to two nascent copies (sister chromatids SC1 and SC2, blue and red). Replication forks (green) then move independently and stochastically in opposite directions along the chain resulting in the growth of the replication bubble. (C) Scheme with all the main steps of the implemented 1D replication dynamics. (D) The two models for sister fork spatial interaction considered in this work.

In this work, we introduce a novel polymer model that combines the simulation of the local 3D chain dynamics with an explicit simulation of 3D chain duplication. Starting from a single origin system, we characterize the effect of a growing replicon (or “replication bubble”) on the structural properties of the chain as function of the speed of the forks along the chain. We then investigate in physiological conditions what is the influence of multiple growing replication bubbles in reshaping the spatial organization of the polymer chain at large scale. Finally, we address the segregation dynamics of the two copy of the DNA segment after full replication and compare some our predictions with experimental data in yeast *Saccharomyces cerevisae*, in order to provide qualitative insights into the nature of replication factories *in vivo*.

## II. RESULTS

### A. A minimal polymer model of genome duplication during replication

To characterize the 3D organization of chromosomes during DNA replication, we introduce a thermodynamic framework that combines a 1D stochastic description of the replication process [27, 28] with a 3D polymer model of the chromosome spatio-temporal dynamics[21], including explicitly chain duplication.

Briefly (see Materials and Methods for a detailed description), a single chromosome is initially modeled as a generic self-avoiding, semi-flexible chain [21] of size *L* (Fig. 1A) evolving under periodic boundary conditions to control the volumic density. Along the chain, a set of monomers is designed as origins of replication and represents the specific regions where replication events may stochastically fire. A replication event initiates with the duplication of the fired origin by adding a new monomer to the simulation and connecting it to the two neighbors of the origin (Fig. 1B, Fig. S1A). This effectively creates two homologous copies of the same monomer embedded into the initial, unreplicated chain. The newly formed structure, termed “replication bubble”, contains two monomers with triple connectivity at its extremities, which represent the so-called “replication forks” (Fig. 1B, Fig. S1A). These forks then stochastically progress in opposite directions along the chain by translating the triple connectivity to adjacent monomers and adding new duplicated monomers to the system at a constant rate *v* (Fig. 1B,C,Fig. S1B). This leads to the growth of the replication bubble. When multiple origins are fired along the chain, the encounter of convergent forks moving in opposite directions results in the merging of the two corresponding consecutive replication bubbles into a larger one (Fig. 1C, Video S1,Fig. S1C,D). At the end of the replication process, the system is made up of two disconnected chains of the same size *L*.

To investigate the impact of the putative pairing between the two forks originating from the same firing event [14, 17, 33] — the so-called “sister forks” — we add the possibility to enforce the spatial co-localization of sister forks during the simulations (Fig. 1 D right), until the corresponding replication bubble dissociates either by merging with another one or by reaching one polymer end. In this case, colocalized sister forks act as one loop-extruder, extruding dynamically two loops — the sister chromatids — as they progress along the chain in opposite directions (Video S2). To mimic a biologically-relevant context, unless specified, we consider in most of our work, the replication of a chain at a physiological speed of *v* = 2.2 kbp/min and comprising *L* = 1000 beads, each of size 25 nm and containing 1250 bp of DNA, at a volume fraction of 6 % before replication, as commonly assumed for *Saccharomyces cerevisae* yeast chromosomes [21](see Materials and Methods).

### 3. Replication bubble in single origin system

To systematically address the role of replication on chromosome organization, we start by investigating a simple system composed of one polymer chain with a single origin localized at its center. The chain conformations are initially equilibrated and, at a given time, the origin is fired and the synthesis of the replication bubble proceeds as described previously. In parallel, we track as a function of time the spatio-temporal dynamics of the polymer during and after replication (Fig. 2A). In particular, we monitor *R*(*N, M*), the spatial distance between two monomers symmetrically located on both side of the origin, separated by a genomic distance *N* and embedded in a structure where *M* monomers have been already replicated. When *M* > *N*, both monomers are located inside the same replication bubble while, when *M* < *N*, they lie on the linear, unreplicated portion of the chain. For example, *R*(*N*, 0) corresponds to their pairwise distance before replication started, i.e., in a linear topology.

**FIG. 2.**
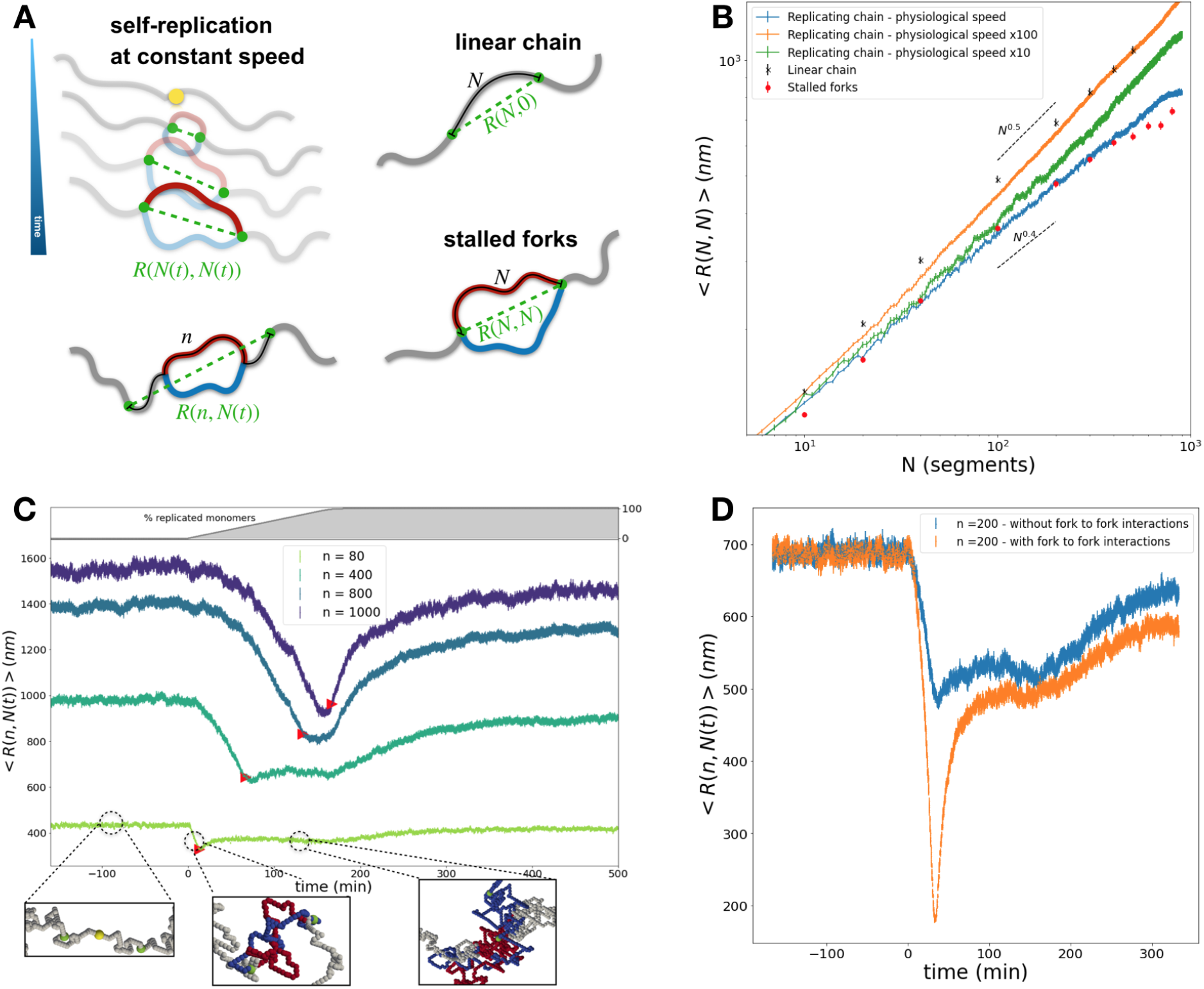
Single origin system. (A) Scheme of the three architectures analyzed in panel B. Green dashed lines indicate examples of distances *R*(*N, M*). (B) Average fork-to-fork 3D distance ⟨*R*(*N* (*t*), *N* (*t*))⟩ as a function of the average number of replicated segments ⟨*N* (*t*)⟩ at various speeds (full lines). For comparison, average 3D distance between *N* segments in a non-replicated linear polymer ⟨*R*(*N*, 0)⟩(black crosses) and between stalled forks ⟨*R*(*N, N*)⟩ (red circles) are given at equilibrium.(C) Time evolution of the average 3D distance ⟨*R*(*n, N* (*t*))⟩ between pairs of symmetrically-located beads separated by *n* monomers (full lines). The red triangles indicate the average timings of replication of both monomers (⟨*N* (*t*)⟩ = *n*). The top panel shows the percentage of replicated chain in time. Three snapshots illustrate typical structures when ⟨*N* (*t*)⟩ ≪ *n* (monomers external to the replication bubble), ⟨*N* (*t*)⟩ ≈ *n* (forks positioned around monomers) and ⟨*N* (*t*)⟩ > *n* (monomers inside the bubble). (D) Comparison for *n* = 200 of ⟨*R*(*n, N* (*t*))⟩ for the non-interacting and interacting scenario.

#### 1. Non-interacting sister forks scenario

First, we consider a model where sister forks do not specifically interact (Fig. 1D left). At a given time *t*, two forks, which are separated by *N* (*t*) replicated monomers along the chain, are then distant by *R*(*N* (*t*), *N* (*t*)) in 3D space. We define ⟨*R*(*N* (*t*), *N* (*t*))⟩ as the population average of the fork-to-fork distance over many simulated trajectories. The progression of forks being stochastic, at a given *t*, each trajectory can have a different *N* (*t*). Therefore, in Fig. 2B, we plot ⟨*R*(*N* (*t*), *N* (*t*))⟩ as a function of the average number of replicated segments ⟨*N* (*t*) ≡ 2(1 + *v t*)⟩.

At physiological replication speed (*v* = 2.2 kbp/min), we observe that ⟨*R*(*N* (*t*), *N* (*t*))⟩ scales as ⟨*N* (*t*) ^0.4^ ⟩(blue line in Fig. 2B). This suggests that the out-of-equilibrium topology, consisting of a polymer ring of growing size connected to two linear branches at the extremities, is more compact that a corresponding linear chain at equilibrium (black crosses, ⟨*R* ∼ *N* ^0.5^)⟩. Such a scaling exponent (0.4) is in agreement with the spatial compaction previously observed in simulations of topologically-constrained ring polymers in semi-dilute conditions at equilibrium [34–37] which was described as a crossover regime towards a universal scaling law with exponent 1*/*3 for larger — crumpled — loops [38]. Consistently, we find that ⟨*R*(*N* (*t*), *N* (*t*))⟩ behaves very similarly to the average 3D distance ⟨*R*(*N, N*)⟩ between stalled forks in a system with replication bubbles of fixed size *N*, at equilibrium (red circles).

Moreover, since the stalled situation corresponds to the limit of infinitely-slow replication, it means that at the physiological replication speed, polymer conformations inside replication bubbles are mostly equilibrated. However, for large bubbles (*N* > 400), ⟨*R*(*N* (*t*), *N* (*t*))⟩ become significantly less compact than the corresponding stalled configuration, suggesting that at large scales, the system remains out-of-equilibrium. This highlights the interplay that exists between the replication time-scale (∼ *N*/*v*) and the typical relaxation time-scale of the replication loop (∼*N* ^2^*τ*_0_ with *τ*_0_ is the characteristic diffusion time of one monomer [39]): for *N* ≲ 1/(*vτ*_0_), bubbles undergoing replication are structurally equilibrated, while for *N* ≳ 1*/*(*vτ*_0_), they remain out-of-equilibrium. In an extreme scenario where the replication speed is very high (*v* = 220 kbp/min, orange line in Fig. 2B), the system does not have the time to relax at any scales and keeps the “memory” of the initial — equilibrated — organization of a linear topology with a scaling exponent of 0.5. In an intermediate regime (*v* = 22 kbp/min, green line), replication bubbles re-equilibrate only at short scale while a crossover towards the linear topology behavior is observed at larger scales.

After the generic characterization of the folding of replication bubbles, we ask how replication dynamically reshapes the relative spatial organization between two loci at physiological replication speed. In particular, we analyze the time-evolution of the 3D distance ⟨*R*(*n, N* (*t*))⟩ between two monomers separated by *n* segments and symmetrically-positioned with respect to the origin (Fig. 2A, bottom left). For every *n* values, we observe that ⟨*R*(*n, N* (*t*))⟩ systematically decreases after the onset of replication (Fig. 2C, Fig. S2A,B), reaching a minimum around the mean replication time of the investigated monomers when sister forks coincide approximately with the monomers’ positions (i.e., when ⟨*N* (*t*)⟩ = *n* ;Fig. 2C, second snapshot). The relative decrease in distance is stronger for larger loops and could be explained by simple scaling arguments: ⟨*R*(*n, N* (*t*) ∼ *n*)⟩ */*⟨*R*(*n*, 0)⟩ ∼ *n*^0.4^/*n*^0.5^ = *n*^−0.1^). After both monomers have been replicated and are within the same replication bubble, ⟨*R*(*n, N* (*t*))⟩ stabilizes to an almost constant distance systematically lower than the initial ⟨*R*(*n*, 0)⟩ value. This intermediate state is abruptly disrupted as one of the chain-ends is replicated and the ring structure vanishes, with ⟨*R*(*n, N* (*t*))⟩ then slowly converging to the corresponding equilibrated distance in a linear topology, but at twice the volume density imposed before the onset of replication. The presence of this intermediate, lower plateau for ⟨*N* (*t*)⟩ > *n* is characteristic of self-avoiding, equilibrated and topologically-constrained crumpled polymer rings, whose internal distances have been shown to be relatively insensitive to the total loop size [36, 37] (Fig. S2C,D).

Overall, our results suggest that the sole process of self-duplication at physiological speed is sufficient to observe a transient decrease of spatial distances between monomers with similar replication time, through the sole effect of entropic forces. Furthermore, this decrease may be partially maintained throughout the replication process.

#### 2. Interacting sister forks scenario

Several experimental studies have suggested that there may exist specific fork-fork interactions that maintain them in close spatial proximity during replication [17]. We test this scenario by including a spring potential between sister forks (see Materials and Methods, Fig. 1D). In particular, we address how, at physiological speed, this new constraint on the bubble’s extremities may affect the entropically-driven compaction process previously described.

As we impose a strong spring constant between the two forks, we find, as expected, a constant value for ⟨*R*(*N* (*t*), *N* (*t*))⟩ regardless of the bubble size (Fig. S3A). Regarding the time evolution of internal distances ⟨*R*(*n, N* (*t*))⟩ (Fig. 2D and Fig. S3B,C) between symmetrically-positioned monomers, we observe the same qualitative behavior than without fork-fork interaction (see above) with a decrease followed by an increase. The minimum distances are still observed around their average replication times, however, it is at much lower values due to the forced co-localization of the forks. Interestingly, as replication progresses and the two monomers exit the region close to the forks (*N* (*t*) > *n*), an intermediate plateau is again recovered. For small genomic distances (*n* ≲ 100), plateau values are very close to the ones observed in the non-interacting fork scenario (Fig. S3C), even if consistently slightly less, due to the presence of two embedding rings, each of size *N* (*t*)/2, that now compose the replication bubble. For larger genomic distances (*n* ≳ 100), the differences between both scenarios are stronger and are maintained over a long time, well after the full replication has been achieved (Fig. 2D and Fig. S3C).

#### 3. In vivo live-cell imaging is consistent with the interacting scenario

In the previous sections, we demonstrate how the replication process can affect the spatial organization of the polymer. While both our considered scenarios for sister forks’ specific interactions lead to a qualitatively-similar increase in compaction within the replication bubble, the magnitude of this effect is found to be stronger under the interacting hypothesis.

In their seminal work, Kitamura *et al*. [17] proposed an experimental set-up where they monitored, in yeast, with live-imaging, the time-evolution of the 3D distance between two loci 60 kbp apart and symmetrically positioned with respect to an isolated origin of replication within chromosome IV along the S-phase (Fig. 3A). In our toy, single-origin system, this procedure amounts to tracking ⟨*R*(*n* = 48 monomers = 60 kbp, *N* (*t*))⟩ as a function of time. Experimentally, they observed a similar dynamics as predicted theoretically by our model, and detected a maximal decrease in distance of about 50 % around the mid-replication time (bars in Fig. 3B). In the paper, the authors considered these observations as proof of the existence of fork-fork interactions. However, we show above that even in absence of such explicit interactions, we may observe a significant decrease in relative distances. Therefore, we quantitatively compare our predictions in both interaction scenarios with their experimental data (full lines, Fig. 3B).

**FIG. 3.**
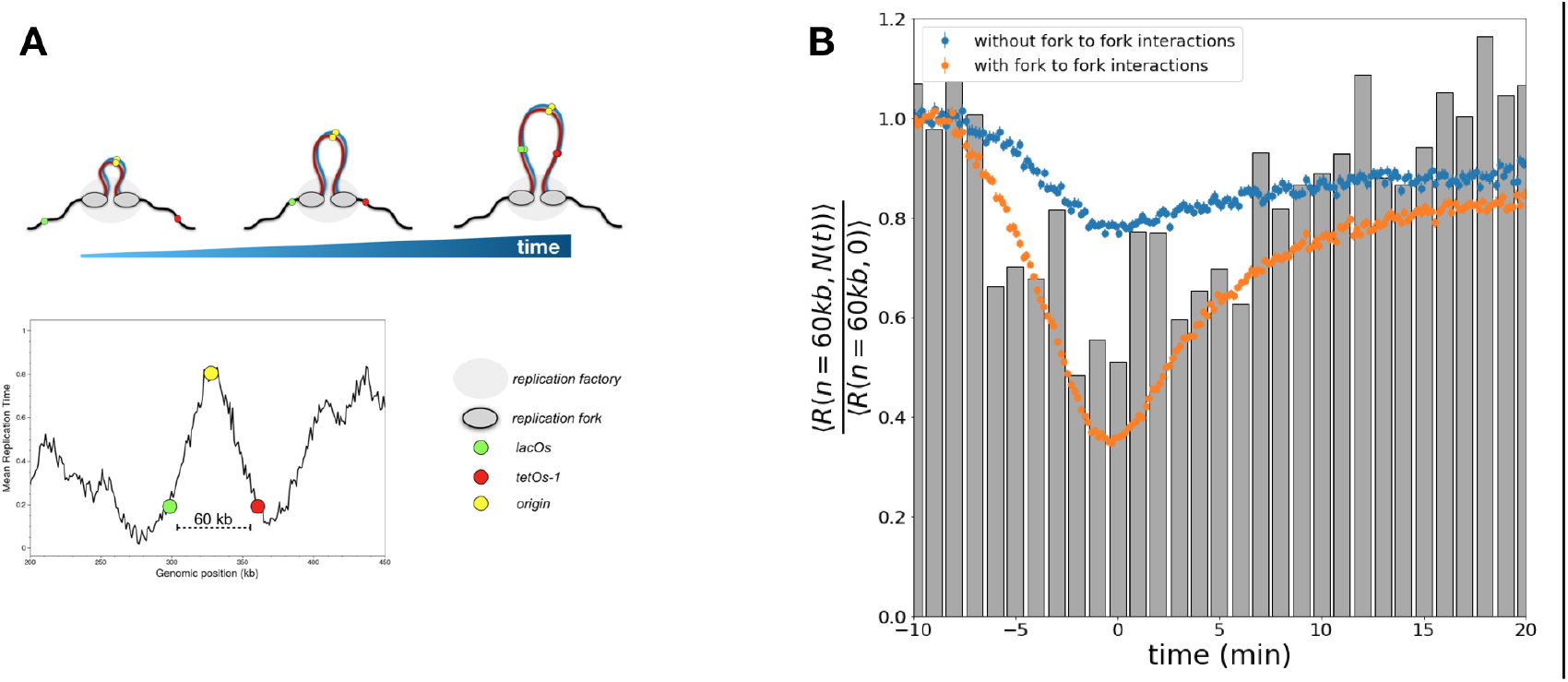
Comparison with *in vivo* live-cell imaging data. (A) Scheme of the experimental setup of Kitamura *et al*. adapted from Figure 4A in [17]. Two chromosomal loci symmetrically positioned around an isolated origin of chromosome IV are tagged with a lacOs and tetOs-1 array. (Top) Progression of the replication forks in presence of putative interacting forks. (Bottom) Experimentally-determined Mean Replication Time profile (MRT) (from [40]) in the region of interest. Higher MRT values correspond to early replicating regions. The yellow circle identifies the origin while the green and red circles indicate the lacOs and tetOs-1 arrays respectively. Given that lack of nearby origins, the two symmetrical loci exhibit similar MRT. (B) Median 3D distance between the two loci normalized by the pre-replication value (experimental values averaged over the first six minutes of acquisition) (grey bars). Corresponding simulated ⟨*R*(*n* = 60 *kb, N* (*t*))⟩ normalized by ⟨*R*(*n* = 60 *kb*, 0)⟩ for both scenarios (full lines).

We confirm that the interacting, loop-extruding hypothesis is more consistent with the available live-cell imaging. Indeed, although the limited statistics of experimental data prevent a more quantitative comparison with simulations, the moderate decrease of ∼ 20% in the case of non-interacting forces seems insufficient to describe the observed degree of co-localization. In contrast, a more substantial decrease of ∼ 65% is observed in the case of interacting forks. Note that this simulated effect is stronger than observed in the experiment, which may be attributed to the constant interaction between forks imposed in the model — while isolated, single forks have also been observed *in vivo* [15]. Under a mixed scenario, this may suggest that about only two third of the replication bubbles have interacting sister forks.

### c. Replication of polymers with multiple origins

Eukaryotic genomes, due to their large linear sizes, necessitate the firing of multiple origins to complete replication in a limited amount of time. To investigate how the simultaneous activity of several replication bubbles may now affect the folding properties of the genome, we simulate a “yeast-like” chromosome with 25 origins regularly inter-spaced along the chain (Video S2). To simplify, we impose the deterministic, simultaneous firing of all the origins at the same time (see Materials and Methods, Video S3). This initial choice of parameters leads to replication timing, maximal bubble sizes and origin density of the same order of magnitude than those observed in yeast [27].

#### 1. Local organization around origins of replication

As shown in the previous single-origin model (Fig. 2D), the presence of a “ring-like” replication bubble can lead to strong local rearrangement of the spatial chain properties as compared to a linear, non-replicating topology. To highlight the structural changes in presence of multiple origins, we compute the time-evolution of the contact probabilities between any pairs of monomers during and after replication in the interacting and non-interacting scenarios (Fig. 4B, Fig. S4 and Video S4). For this computation, to avoid ambiguity due to the incomplete replication of the polymer, we consider only contacts between monomers belonging to the same chromatid, between still-unreplicated monomers or between a monomer on one chromatid and another one still unreplicated (see Materials and Methods). For both scenarios, Figure 4A shows the resulting contact maps 3 minutes after replication started, averaged around an origin and normalized by the corresponding map in absence of replication. Full time course along the S-phase can be found in Fig. S4A,C (see also Video S4). Whatever the scenario, we observe that replication dynamically impacts the contact patterns. In the non-interacting fork context, the previously-discussed compaction due to the topological property of polymer rings, is in fact sufficient to observe a weak, but significant, local increase in contacts around origins and inside the bubble (label 1 in Fig. 4A,B and Fig. S4A). Under the interacting-fork hypothesis, we observe the distinct signature of a loop between the current positions of two sister forks. As the replication proceeds bidirectionally and symmetrically, the locations of this loop move perpendicular to the main diagonal of the contact map (Fig. S4C,D). The paired sister forks are thus equivalent to a bidirectional loop extruder, loaded at the origin.

**FIG. 4.**
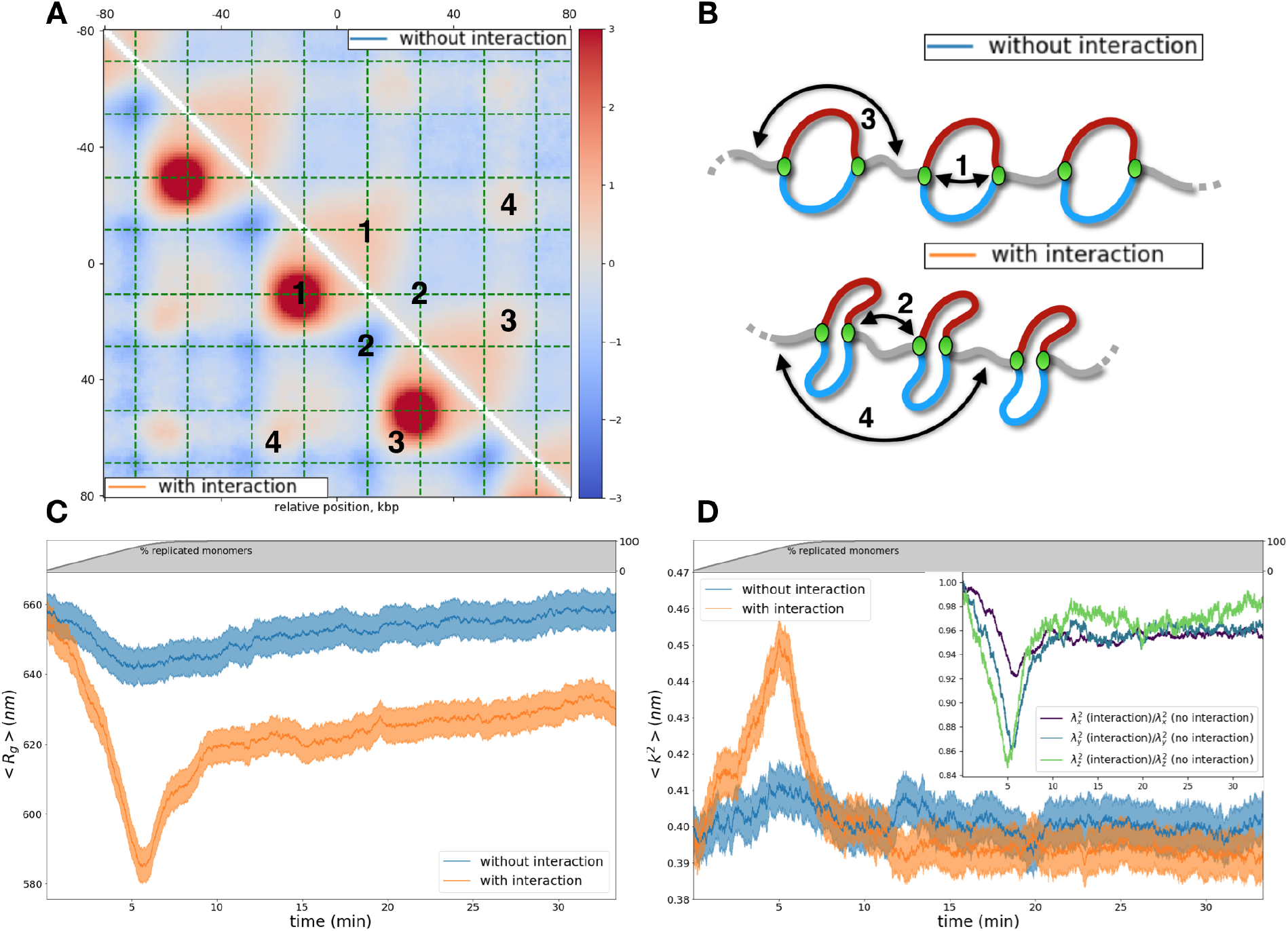
Structural properties of multiple origin systems. (A) Average normalized contact map around an origin of replication 3 minutes after replication have started. The upper and lower triangles show the non-interacting and interacting conditions, respectively. Red(ish) (resp. blue(ish)) colors mean more (resp. less) contact than in the linear - unreplicated - topology. Dashed green lines indicate the average fork positions at the specific time step. Numbers on the maps are specific contacts depicted in panel B. (B) Scheme summarizing some principal elements of the system: contacts between sister forks (1), convergent forks (2), linkers separated by 1 bubble (3), linkers separated by 2 bubbles (4). (C,D) Time-evolution of the average radius of gyration ⟨*R*_*g*_ (*t*)⟩ for interacting and non-interacting forks (C) and of the relative shape anisotropy ⟨*k*^2^(*t*)⟩ for interacting and non-interacting forks (D). (Top) Percentage of replicated monomers as a function of time. (D, Inset) Ratio for each component of the gyration tensor between the two conditions.

Additionally, in both cases, the ongoing compaction of individual replication bubbles results in an increase of contacts between distal linker regions, i.e., between the unreplicated parts of the chain separating two convergent forks (labels 3 and 4 in Fig. 4A,B). Interestingly, in contrast, a decrease of contacts is observed between consecutive convergent forks (label 2 in Fig. 4A,B). Such an effect is much stronger in the interacting scenario and may be due to entropic repulsion between two consecutive bubbles (see also below) that may ‘extend’ the linker.

#### 2. In vivo 3C data are consistent with an extrusion-like scenario

These results highlight how replication may lead to non-trivial features on contact maps. In particular, in the context of interacting — loop-extruding — forks, loops can be clearly seen. We thus wonder whether such a signature can also be observed in experimental data.

While chromosome conformation capture (3C) techniques can now give access to genome-wide contact frequencies between any pairs of genomic loci, addressing experimentally the time evolution of 3D chromosome organization during S-phase remains challenging. However, for the yeast *Saccharomyces cerevisiae*, such data are now available [5, 13]. We first processed Micro-C (a high-throughput, whole-genome 3C method) data from Costantino *et al*. [13], in which the authors synchronized cells in G1 and then measured a time course of the 3C contacts at different times after the release. To be comparable with our predictions, we compute the normalized average Micro-C contact maps around early replicating origins [40] (see Materials and Methods) (Fig. 5A and Fig. S5A,S6A). Remarkably, we observe a strong enrichment perpendicular to the main diagonal on a short range (10 − 20 kb) in early S phase, which extends further in late S and gets gradually weaker in S/G2 and G2 phases (Fig. 5A). Such a signature is not visible around late-replicating origins (Fig. S5C) suggesting that it is specific to loci that are frequently and synchronously fired in early S-phase.

**FIG. 5.**
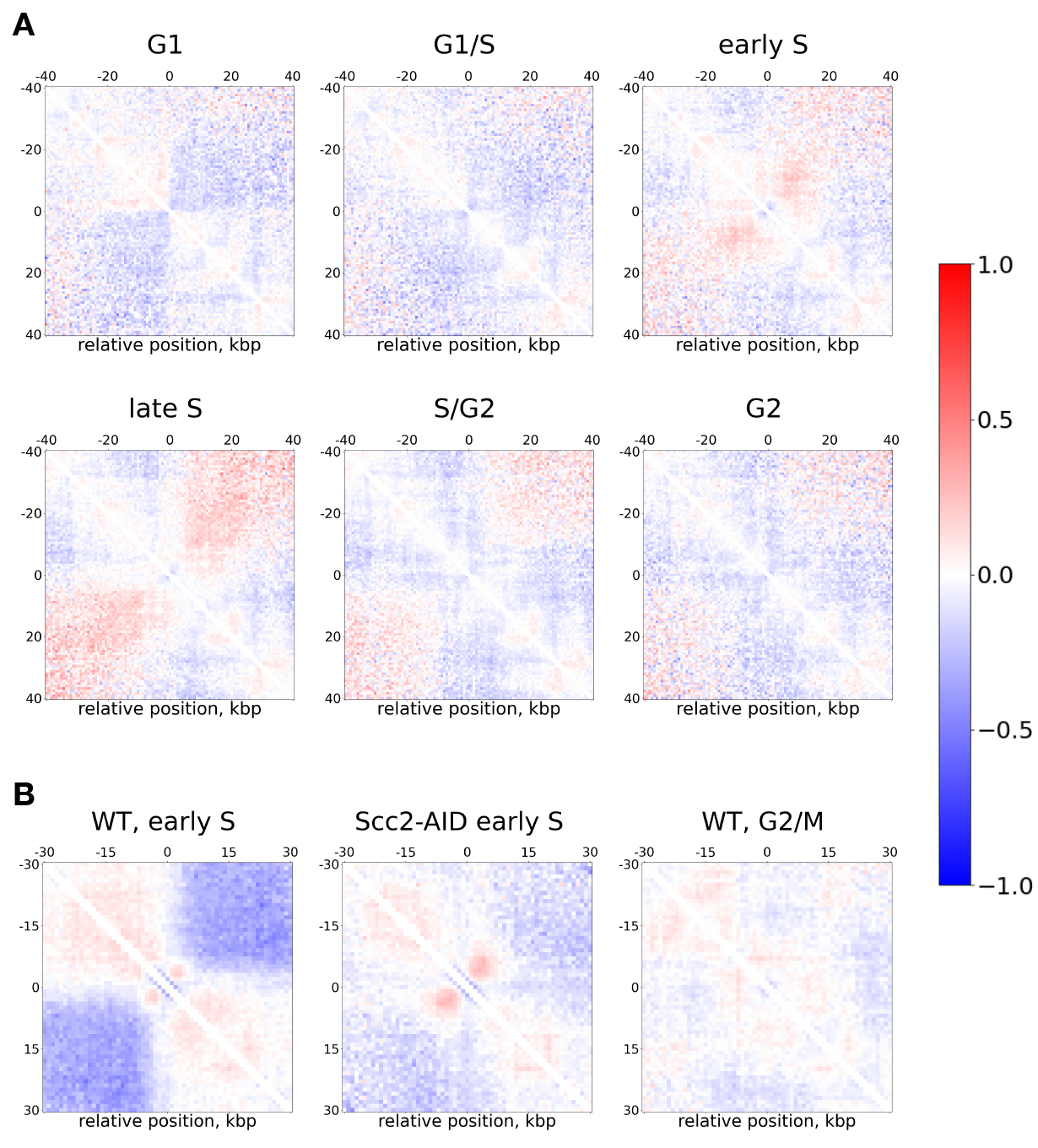
Experimental contact maps around strong origins of replication in *Saccharomyces cerevisiae* . (A) Average normalized (Observed over Expected) contact maps at 800 bp resolution around early replicating origins using experimental data from [13] at different stages of the cell cycle. (B) As in (A) but using experimental data from [42] at 1 kbp resolution. The three plots show three different experimental conditions: Wild type (WT) cells arrested in Early S with HU treatment, scc2 depleted cells arrested in Early S with HU treatment and G2/M arrested cells.

However, during S-phase in yeast, cohesin complexes, also capable of performing DNA loop extrusion, [5, 13, 41] are concomitantly loaded onto chromatin. Indeed, the original conclusion by Costantino *et al*. from the analysis of the same dataset was the progressive accumulation of chromatin loops between cohesin associated regions (CARs), the genomic loci enriched in cohesins, during the S-phase. This introduces some potential ambiguity in deciphering the role of fork-mediated extrusion in contact maps, as we cannot exclude that the short range enrichment in early S-phase may be also caused by cohesin-mediated extrusion. To partially address this potential bias, we perform several complementary analyses (see Materials and Methods). First, we compute the average Micro-C contact maps around early-replicating CARs (Fig. S5D,E). As observed in [13], cross-like patterns, very different from the motif seen around early origins, appear during S-phase and are maintained during G2. They are the signature of CAR-CAR loop mediated by cohesin. Then, we estimate the average cohesin ChIP-seq signal around early-replicating origins in early S (Fig. S5F,G). We do not observe specific enrichment that might have been expected if the observed pattern around origins were driven by cohesin. Finally, we analyze two other yeast datasets [42, 43], where contact frequencies in early S-phase were measured. In the first one [42], cells were arrested in very early S (using hydroxyurea [HU] treatment) in the presence (WT) or absence (Scc2-depleted) of cohesins (Fig. 5B,C and Fig. S6B). The original objective of this study was to investigate the role of replication forks in stalling cohesin-mediated loop extrusion. Using this dataset, we again observe a clear enrichment of contacts perpendicular to the main diagonal around early origins for the cohesin-depleted cells and also, but at a lesser extent, in WT condition. The lower signal in WT suggests a competition beween cohesin and replication for the shaping of the 3D structure of the chromatin. Also in this case, no clear enrichment is detectable anymore in G2/M. In the second dataset [43], cells were ‘synchronized’ in S-phase with a milder and longer HU treatment. In this case, we do observe a fountain-like signal but only around early-replicating origins far from centromeres (> 40 kbp) (Fig. S7). Overall, our analysis corroborates the presence of a cohesin-independent contact signature around early origins that resembles the predicted pattern generated in the replication-dependent interacting - extruding - scenario. Note, however, that the strength of contact enrichment observed experimentally is weaker than the predicted one. This may be due, as for the *in vivo* live imaging, to a mixed scenario where only a fraction of the sister forks remains attached and thus effectively performing loop extrusion. Furthermore, experimental data are likely more heterogeneous that our “ideal” system, for which the observed patterns are clearly magnified by the symmetry and perfect synchronization of the origins.

#### 3. Impact of loop extruding sister forks on the large-scale organization

Beyond the local reorganization observed in contact maps during replication, we then address how such local conformational changes may impact the large-scale folding of the polymer chain.

To characterize the global size of the chain, we track the time-evolution of the average radius of gyration ⟨ *R*_*g*_(*t*)⟩ (see Materials and Methods) during and after replication in both sister fork scenarios (Fig. 4C). Like for the internal distances within replication bubbles (Fig. 2D), we observe a dynamic decrease of ⟨*R*_*g*_(*t*)⟩ after replication has initiated, which is more marked in presence of fork-fork interactions (Fig. 4C). The global compaction reaches a maximum value (minimum ⟨*R*_*g*_(*t*))⟩ as individual replication bubbles reach their maximal sizes just prior to merging events between convergent forks and the replication of chain ends. Such an effect is more pronounced in the loop-extruding (interacting forks) scenario compared to non-interacting forks (∼ 12% vs ∼ 3% decrease). In the latter case, the overall structure quickly relax within 20 minutes to its initial compaction value. However, in the former, the slow relaxation dynamics maintain the chain in a more compact state for a very long time (see also below).

In addition to ⟨*R*_*g*_(*t*)⟩, we also monitor the relative shape anisotropy ⟨*k*^2^(*t*)⟩ of the whole chain (see Materials and Methods) (Fig. 4D): *k*^2^, for a cloud of points, quantifies its degree of anisotropy with *k*^2^ ≈ 0 for spherical arrangements and *k*^2^ ≈ 1 for rod-like shapes. Only in the interacting, extruding scenario, we remark a significant change in anisotropy. The polymer becomes more elongated as replication proceeds. The peak in anisotropy coincides with the timing of maximal compaction observed in Fig. 4C. The transition towards a more cylindrical structure can be further characterized by comparing the individual components of the gyration tensor (Inset in Fig. 4D) with a significant decrease of the second (12.5 %) and third (15 %) components. Merging events at the end of replication abruptly destabilize such a structure with a fast relaxation towards values close to the initial ones.

All these results demonstrate how the local ring topology imposed by sister-fork interactions reverberates on the global organization of the polymer. In particular, the loop extrusion activity in the interacting scenario drives transiently the system close to a “polymer bottle-brush”-like architecture [44–47] for which the presence of multiple, consecutive, large replication bubbles (Fig. 4B, bottom) enhance the loop-loop entropic repulsion and lead to a relative elongation of the structure. This effect is reminiscent of polymers with dense arrays of consecutive loops and is associated to chromosome condensation via SMC-mediated loop extrusion [44–47]. In the context of replication, density and size of loops are controlled by the origin density and replication timing that may locally limit the bottle-brush effect in a specific time-window where replication bubble lengths are maximal.

### D. Relative organization and replication-driven catenation of the sister chromatids

In the previous sections, we focused on the internal organization of the replicating chain. Here, we finally characterize the relative organization of the sister chromatids after full replication.

For this purpose, we start by investigating how the distance between the centers of mass of each sister chromatid ⟨|*R*_*cm*2_ − *R*_*cm*1_ (*t*)|⟩ evolves as a function of time for the two sister forks interacting scenarios and for a system with multiple (25) origins, and we compare with simulations at equilibrium of two separated chains of the same size evolving at similar density (Fig. 6A). Overall, we observe a slow gradual increase of the relative distance as a function of time, likely reflecting the ongoing individualization of each chromatid. No clear distinction is visible between the two scenarios and the relative distance grows with a sub-diffusive behavior (⟨|*R*_*cm*2_ − *R*_*cm*1_ |(*t*)⟩ ∼ *t*^0.3^) and seems to very slowly converges towards the equilibrium value obtained when the two chain starts at separated positions. This suggests that sister chromatids after replication have a constrained relative motion that may be the signature of the intertwining between the two sister chromatids, observed in simulation snapshots just after replication (Video S5 and Fig. 6B), that may hinder the diffusion of individual chromatids. Indeed, in a system where the two chains are initially fully-overlapping (corresponding to a situation of instantaneous replication with every monomer being an origin, see Materials and Methods) - thus leading to a high degree of intertwining (Video S5) - we observe similar dynamics (red line in Fig. 6A), consistently with simulations of segregating chains performed by Dockhorn *et al*. [48].

**FIG. 6.**
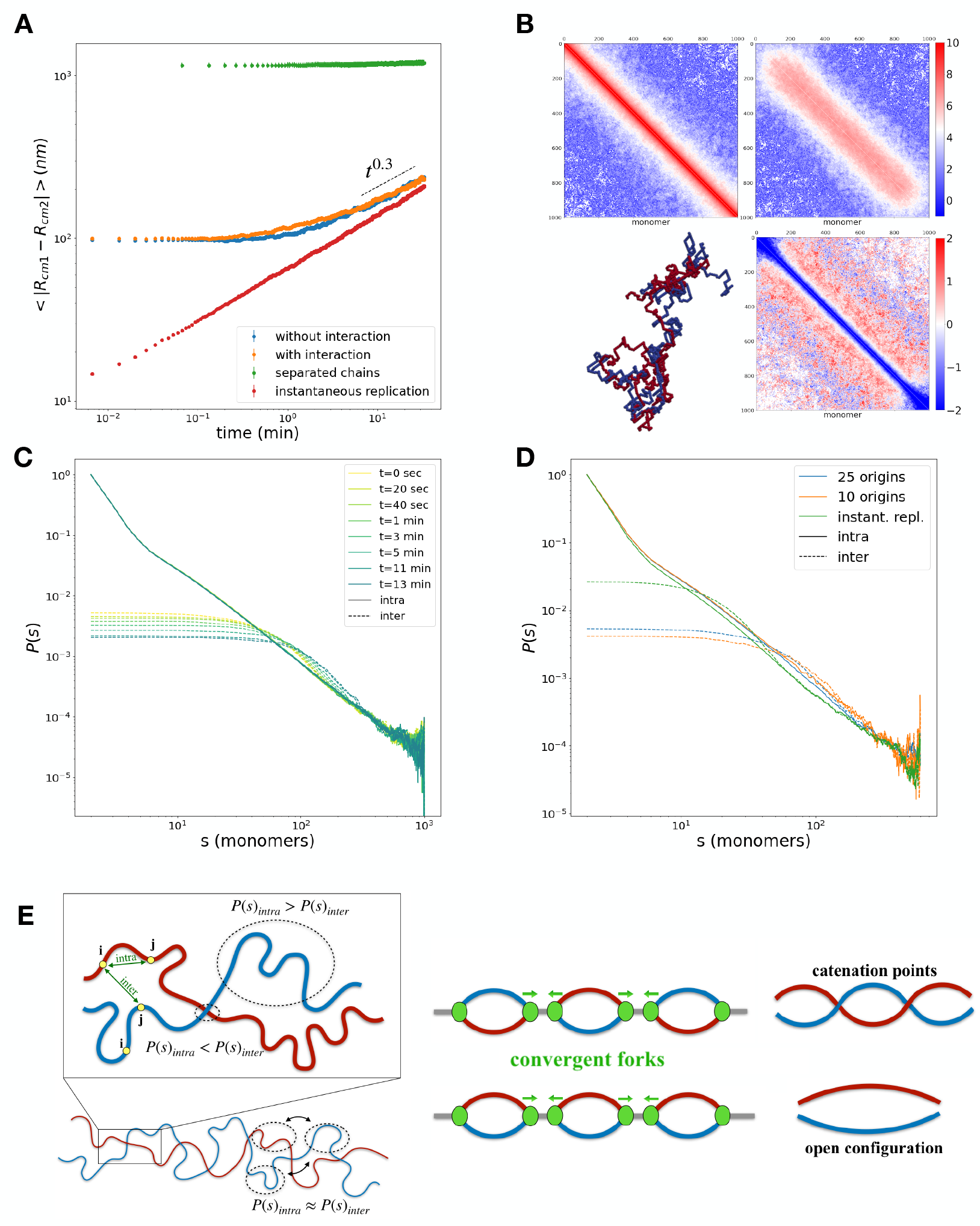
Relative organization of the sister chromatids. (A) Time evolution of the distance between the centers of mass of the sister chromatids ⟨|*R*_*cm*2_ − *R*_*cm*1_|(*t*)⟩ for a system with 25 origins of replication in the non-interacting forks (blue) and interacting forks (orange) scenarios, following full replication. The green line corresponds to the equilibrium values for two chains and the red line to two chains created by instantaneous replication. (B) (Top left) Intra-contact map and (Top right) inter-contact map at time= 13 min after full replication. (Bottom right) Log-ratio between the inter-contact and intra-contact maps shown above. (Bottom left) Snapshot of a simulation at the same time illustrates the intertwined sister chromatids with loose ends. (C) Average intra (solid curves) and inter(dashed curves) contact probability as a function of the linear distance *s* along the chain for different time values *t* after full replication, for a system with 25 origins of replication with sister-forks interactions. (D) As in (C) but at *t* = 0 after replication for three different origin densities and in absence of sister-forks interactions. (E) Scheme to illustrate the spatial organization of sister chromatids following replication.

To characterize further how this post-replicative intertwined state — which is independent of the sister-fork scenario (Fig. S8) — evolves in time, we measure 3C-like maps of intra-chromatid and inter-chromatid contacts (see Materials and Methods) after the end of the replication process (Fig. 6B, top). As the corresponding matrices are mostly invariant along their sub-diagonals (except around the chain ends), we compute the average value *P*_*intra*_(*s*) and *P*_*inter*_(*s*) along each sub-diagonal *s* (Fig. 6C,E, Fig. S8). *P*_*intra*_(*s*) corresponds to the average contact probability between any two monomers *i* and *j* belonging to the same chromatid and separated by a linear distance *s* =|*j* − *i*| along the chain. Similarly, *P*_*inter*_(*s*) is the average contact probability when *i* and *j* are on separate chromatids. For example, if both sister chromatids are perfectly aligned, we expect *P*_*intra*_(*s*) = *P*_*inter*_(*s*).

*P*_*intra*_(*s*) scales as *s*^−3/2^ and is independent of time, suggesting that intra-chain contacts are controlled by chain connectivity and behaves as an equilibrated polymer [49]. For *P*_*inter*_(*s*), at each time step *t*, we observe three regimes (Fig. 6 C,E): (1) for *s* < *s*_*min*_(*t*), *P*_*inter*_(*s*) is almost constant and *P*_*intra*_(*s*) > *P*_*inter*_(*s*) ; (2) for *s*_*min*_(*t*) < *s* < *s*_*max*_(*t*), *P*_*inter*_(*s*) exhibits a “shoulder” with *P*_*intra*_(*s*) < *P*_*inter*_(*s*) ; (3) for *s* > *s*_*max*_(*t*), *P*_*inter*_(*s*) ≈ *P*_*intra*_(*s*) ∼*s*^−3/2^. *s*_*min*_(*t*) (≈ 40 monomers at *t* = 0 after replication and ≈ 60 at *t* = 13 min) and *s*_*max*_(*t*) (≈ 200 monomers at *t* = 0 and ≈ 300 at *t* = 13 min) are both increasing as a function of time. The first regime corresponds to a scale where both chromatids are largely segregated. The second regime is mainly driven by the catenation between distal points on the two separate chromatids. Finally, in the third regime, beyond the typical catenation scale, both chromatids are maintained together by the various catenations along the chains and are loosely aligned, thus leading to similar contact probabilities.

The dynamic evolution of *s*_*min*_(*t*) and *s*_*max*_(*t*) shows that the chromatids become locally more and more segregated and that the linear distance between the catenation points increase in average as a function of time while both chains relax towards equilibrated — more separated — configurations. Interestingly, the wider area observed in the log-ratio matrix between the inter- and intra-contact maps (Fig. 6B, Fig. S9,S10, Video S6) at the chain ends indicates the more efficient relaxation and segregation of the two chromatids at their freely-diffusing, non-catenated extremities.

To confirm that the observed behaviors are due to topological constraints between sister chromatids, we perform simulations in the limit of instantaneous replication where we allow strand passing while conserving steric hindrance (see Materials and Methods). In this case, we observe a relatively faster diffusion between the centers of mass (⟨|*R*_*cm*2_ − *R*_*cm*1_|(*t*)⟩ ∼ *t*^0.5^)(Fig. S11 A) and the disappearance of the shoulder (*s*_*min*_ = *s*_*max*_ ∀*t*) (Fig. S11).

We then speculate that the main cause of this catenation is the merging events between consecutive replication bubbles emanating from multiple origins due to the random orientation of each bubble (Fig. 6E). To test this hypothesis, we investigate the role of the number of origins in creating such intertwined structures. We proceed with the same analyses for 10 origins and in the limit of instantaneous replication (∼ 1000 origins) (Fig. 6D and Fig. S9C, S10). *P* (*s*) curves just after replication show how decreasing the number of origins reduces the intertwining levels. In particular, fewer origins lead to an increase in both *s*_*min*_ and *s*_*max*_, suggesting less catenation. Notably, in the limit of only one origin (Fig. S8C), *s*_*min*_ = *s*_*max*_ (*P*_*inter*_(*s*) ≤ *P*_*intra*_(*s*), ∀*s*) and there is no more catenation. Conversely, we study the effect of increasing the chain length *L* while keeping the linear density of origins (one origin every 40 monomers) and the chain volumic density constant. Repeating the analysis for *L* = 2000, 3000 and 5000 (see Materials and Methods), we find that the previously described features are qualitatively recovered and do not depend strongly on *L* (Fig. S12), suggesting that the relative dynamics depends on the density of catenations rather than the actual total number of catenations. Quantitatively, however, we observe a systematic slightly higher individualization at small *s* (< *s*_*min*_(*t*), first regime) for longer *L*, which may originate from the gradual increase of topological effects [50].

## III. DISCUSSION AND CONCLUSION

In this paper, we present an original polymer framework designed to investigate eukaryotic DNA replication in 3D at the kilo-basepairs scale. Our minimal model of the replication process enabled us to characterize the physics of a self-replicating polymer, emphasizing the core differences from its linear counterpart. We systematically analyzed the fundamental properties of replication bubbles and sister chromatids in single and multi-origin systems and in presence or absence of interactions between sister forks, demonstrating how the out-of-equilibrium process of replication can lead to significant perturbations in 3D spatial organization. In particular, we showed that the newly formed structures, in physiological conditions, are better described by the physics of polymer rings and loose polymer bottle-brushes rather than that of linear chains. The dynamic transition from linear to ring polymer is responsible for a significant compaction at the local replication bubble level, while also affecting the global conformation beyond the replication forks. Interestingly, a partial structural memory of such 3D reorganization could be stably maintained well after the chain has been fully replicated.

Our model allowed us to make predictions that distinguish the two interacting scenarios for sister forks using measures that are experimentally accessible. Specifically, we demonstrated how the hypothesis involving interacting sister forks exhibits better agreement with both microscopy and chromosome conformation capture data in yeast [13, 17, 42]. In particular, in this scenario, the complexes formed by the two linked forks act as effective loop extruders loaded at specific sites, the origins of replications. As a signature, this leads to a very specific pattern of pronounced enrichment of contacts around origins (Fig. 4A) that are visible on experimental 3C data (Fig. 5). Such spatial features, observed here in the replication context in yeast, are analogous to the ones arising from the direct loop extrusion process mediated by Structural Maintenance of Chromosomes (SMC) complexes like cohesins and condensins when preferentially recruited at distinct loci, and termed as chromatin “fountains” or “jets” in the recent literature [51–53]. Remarkably, a recent experiment using a customized Hi-C-like technique able to capture contacts of early replicating regions also observed fountain-like patterns around origins of replication in human cells [54]. However, this analogy with SMC-mediated extrusion carries also some ambiguity as loop extruding cohesins are also loaded on chromatin during S-phase concomitantly of replication in yeast, or are already present on chromatin since G1-phase in mammals [5, 13, 32, 43]. Although our analysis of existing datasets in yeast suggests that observed fountains around origins may be cohesin-independent[13, 42], future Hi-C experiments — for example in a synchronized population of cohesin-depleted cells — should be performed at different timepoints during S phase to confirm this. To explore if similar looped or fountain-like patterns may emerge through different replication-dependent effective mechanisms than the interacting scenario, we test two alternative models (See Materials and Methods): one where newly replicated monomers can self-attract (Fig. S13) (putatively via the transient binding of specific proteins capable of oligomerizing) and another where the stiffness of the new sister chromatids is different from the mother (Fig. S14) (putatively via the transient depletion of the histone content on chromatin). In both cases, we observe non-trivial patterns around origins but they mostly resemble the contact enrichments seen in the non-interacting case and clearly differ from those observed experimentally and in the interacting-forks scenario. Therefore, even if we cannot rule out the possibility of other mechanisms driving genome folding around origins, our work emphasizes the putative role of replication-dependent, transient loops and effective loop extrusion in genome organization during S-phase and beyond. This may represent a complementary mechanism to SMC-mediated loop extrusion to regulate the local and global compaction and elongation of chromosomes [44–47]. The exact molecular mechanism driving effective fork-fork interactions is not known. Ctf4 may be involved by bridging the two helicases present at sister forks [55]. But any multivalent or oligomerizing molecules that have specific affinity to fork elements or to the chromatin located just nearby forks may potentially lead to such a replication-dependent loop extruding mechanism [56].

As the principal scope of DNA replication is the transmission of genetic information through cell division, characterizing how two copies of the same chromosome interact in space following their duplication constitutes a crucial task. In eukaryotes, the reciprocal positioning of the two sister chromatids in G2 is in part insured by a process called cohesion, also mediated by cohesin complexes that bridge the two sisters regularly along the genome [57–61], that is still poorly understood. Such a relative organization between sister chromatids is instrumental in DNA repair and eventual segregation during mitosis. Our results (Fig.6) suggest that the sole process of replication might drive the system to a preliminary state where the two chromatids maintain a certain degree of cohesion due to chain catenations and intertwinings, without the need for specific bridgers between the two chains, and leads to slow, constrained dynamics between the sister chromatids. In particular, we predicted a very specific dynamic signature of such a cohesion in the intra- and inter-chromatids contact maps (Fig. 6 B). It would be intriguing to compare this with data in early G2, obtained from advanced 3C-like techniques, like Sister-C [57], that allow to separate the 3C-contacts between sister chromatids, which is currently very challenging experimentally and only available for arrested M-phase cells in yeast. Such experiments must be carried out in cohesin-depleted cells, in order to isolate the effects of replication-driven catenation from cohesin-mediated cohesion and loop extrusion and mirror the conditions here simulated.

This replication-dependent catenation — cohesive — effect contrasts with the situation in prokaryotes, where the replication of circular genome from one origin by self-interacting sister forks has actually been theoretically suggested to facilitate sister chromatids segregation due to loop-mediated entropic forces[31].

To conclude, our model offers a quantitative framework to investigate the role of out-of-equilibrium replication in genome structure and dynamics. In this work, we focused on the fundamental polymeric properties of replicating chains, and therefore made several assumptions to simplify the analysis. For example, we mostly simulate chromatin as a topologically-constrained, self-avoiding walk, excluding chain crossing and thus possible effects of Topoisomerases that may act *in vivo*. While we highlighted the role of chain crossing in the relative catenation of sister chromatids, experiments where the Topoisomerase activity can be controlled [62], in cohesin-depleted conditions (see above) to remove putative biases from cohesion, would be required to finely describe quantitatively such an effect. However, our framework is modular and offers the possibilities to integrate various biologically-relevant ingredients and to refine and contextualize our generic conclusions to specific biological systems by implementing more realistic 1D replication dynamics and probing their 3D consequences. In particular, it would be interesting to characterize the interplay between the replication-dependent loop-extrusion-like process described here with other main mechanisms organizing the genome like SMC-mediated loop extrusion, (micro)phase-separation mediated by architectural proteins or localisation at the nuclear periphery, and between replication-dependent catenation and cohesin-mediated cohesion [63, 64]. For example, in yeast, chromosomes follow a Rabl organization with all the centromeres clustered on one side of the nucleus and the telomeres tethered to the nuclear envelope. The emerging structural features predicted by our model are mostly relevant at the intra-chromosomal scale and, predominantly, at the replication bubble level. Therefore, we expect our predictions to be qualitatively valid in the Rabl context. However, the polymer bottle-brush effects emerging from fork-fork interactions may participate in the increase of intra-chromosome contacts (vs interchromosome) observed in S/G2 [65]. Similarly in higher eukaryotes having longer chromosomes (Fig. S12), we do not expect a qualitative modification of the model outcomes. Although, quantitatively, the heterogeneity in chromatin volumic density within the nucleus may lead to differential effects between eu- and heterochromatic regions, the former being generally less dense [66] and thus predicted to be more sensitive to the replication-dependent polymer bottle-brush effect (Fig. S15).

In addition, while here we focused on interactions between only sister forks, it may be interesting to investigate the consequences of attractive interactions between any active forks to address the putative formation of higher-order replication “factories” grouping several replicons, whose existence still remains highly controversial [15, 16]. Finally, our model, while explicitly describing the impact of replication in genome folding, currently lacks a feedback mechanism between the 3D organization and the underlying 1D stochastic dynamics of DNA replication. It could be intriguing to explore the influence of pre-existing 3D organization (driven by other mechanisms described above) and of the replication-dependent 3D changes on the complex temporal pattern of origin activations, for example via the 3D diffusion of a limited number of firing factors that may activate origins differentially depending on their 3D environment [27, 67].

## IV. MATERIALS AND METHODS

### A. Model and simulations

#### 1. Null polymer model

Chromatin is described as a self avoiding walk composed of *L* beads and dynamically evolving on a SxSxS fcc lattice with periodic boundary conditions via a Kinetic Monte Carlo (KMC) algorithm, as previously described [21, 68]. Briefly, each Monte Carlo time step (MCS) consists in *N* trial moves (with *N* the current number of monomer in the system): in a trial move, a randomly-picked monomer is attempted to move to a randomly-chosen neighboring site among those which preserve the connectivity of the chain. To account for self-avoidance, moves are automatically rejected if the trial site is already occupied by a monomer which is not consecutive of the moving monomer along the chain. In addition to excluded volume, the polymeric chain is subjected to a standard potential to account for the chain bending rigidity:

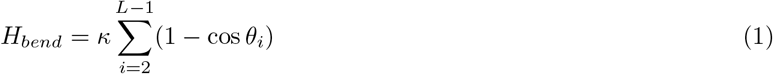

with *θ*_*i*_ being the angle between monomers *i* − 1, *i* and *i* + 1 and *κ* the bending modulus in kT units. The trial move is finally accepted according to the Metropolis criterion on *H*_*bend*_ [69].

*L, S* and *κ* were set to model the chromatin fiber in *Saccharomyces cerevisiae*. Assuming standard values for the chromatin fiber diameter (*σ* = 25 nm) and linear compaction (50 bp/nm) [70], we recover an approximate bead size of 1250 bp and fix *L* = 1001 in order to describe larger yeast chromosomes (∼ 1 Mbp). *S* = 16, assuming a typical base-pair density of 0.007 bp/nm^3^ (6% volumic fraction initially) in yeast nuclei. *κ* = 2.65 kT, assuming a Kuhn length for chromatin of 100 nm [21, 68, 71].

Similarly to [21, 72, 73], time mapping between the simulation MCS and real time is done by computing the Mean Squared Displacement (MSD) of a monomer. We then compared it to the experimental MSD_*exp*_(*τ*) [*µm*^2^]= ⟨((*r*(*t* + *τ*) − *r*(*t*))^2^⟩ ≈ 0.01 · *τ* ^1/2^ with *τ* in seconds, measured in yeast [74], leading to 1 MCS = 0.2 msec.

The separated chains simulations showed in Fig. 6 A were implemented by populating the lattice with two distinct chains of 1000 monomers. The two backbones used to initialize the polymers were located at a distance of *S/*2, where *S* = 16 is the size of the box. The two chains are introduced sequentially into the lattice, taking care that during the growing algorithm the second chain does not cross the one already initialized.

Since the characteristic spatial and time scales of the phenomena under study (∼100s nm, ∼ min) are well beyond the discretization scales (10s nm, ∼msec) imposed by the lattice and the KMC algorithm, the obtained results are not expected to depend qualitatively on the underlying modeling and simulation framework [75].

#### 2. Self-replicating polymer model

To account for the replication of the chain, we complement the previous polymer mode by specialized classes of monomers. A specific set of origins of replication is defined along the chain. At a predefined time, all the origins are instantaneously and deterministically fired. Note that the model can be easily generalized to a heterogeneous set of asynchronized origins with different firing probabilities [27, 40]. Before the onset of replication, every bead is initialized with an integer parameter *status* = 0, assigning it to the template “maternal” chain (Fig. 1A and Video S1). Origin firing consists in the addition of a new monomer in the system, at the exact same position of the fired origin (i.e. on top of the origin). The fired origin and the new added monomer are then connected by two bonds to the two origin nearest-neighbors along the chain and their status are set to − 1 and 1 respectively in order to label the two distinct chain replicates. Due to its biological analogous, the resulting structure is referred as a replication bubble (Fig. 1B, Fig. S1A and Video S1).

The extremities of replication bubbles are termed as forks due to their specific property of being connected to three other beads. The two forks of the same bubble are named sister forks. Each fork is characterized by a directionality according to its orientation with respect to the origin. During the following MCS, every fork in the system can trigger further synthesis with a probability *v* by introducing additional monomers at the fork positions and moving the triple connectivity according to its directionality (Fig. 1B, Fig. S1B and Video S1). Note that, temporarily, the addition of new monomers breaks the excluded volume rule as they are initially positioned on the same lattice node of another monomer. However, as they move afterwards, excluded volume constraint apply again leading such a double occupancy to very rapidly vanish. Since the forks are moving much slower than the local diffusion time of monomers, we do not expect such an effect and more generally the exact chosen strategy to integrate new monomers to have a significant impact on the results as the polymer will locally equilibrate before an other monomer will be replicated.

Note also that the size of the simulation box remains constant during the replication process, meaning that the volumic fraction is doubling between the start and end of the replication, while, *in vivo*, the nuclear volume increases throughout the S-phase [76]. However, at the low yeast-like concentration used in this paper, we do not expect major qualitative differences in the presented results.

In case of multiples origins, two converging replication bubbles merge as a result of the encounter of two forks of opposite directionalities. When two opposite forks converge and are now neighbors along the chain, each one has a reduced probability to replicate equal to 0.5*v*. In fact, when one of the two forks is selected for a replication move, it will automatically replicate also its neighbor, merging the two converging bubbles (Fig. S1C,D). This ensures that the replication speed *v* remains constant also in the case the merging events where the DNA replicated is effectively doubled. An analogous reduced probability rule is applied when a fork encounters a chain end, the bubble is then opened by replicating both the fork and the chain end monomers.

Colocalization of sister forks in the interacting - loop extruding - scenario is maintained by adding an elastic-like potential to the Hamiltonian of the system between the monomers where the forks are at a given time :

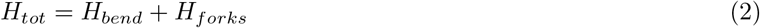

With

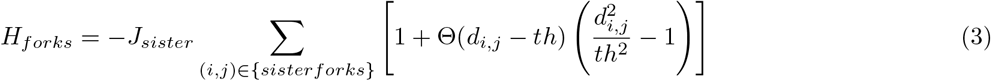

where the sum is performed over all the pairs of still active sister forks localized at monomers *i* and *j, d*_*i*,*j*_ is the 3D Euclidean distance between *i* and *j*, Θ(*x*) is the Heaviside step function and *th* = 50 nm is the distance below which the energy saturates to *J*_*sister*_ = 100 kT. Springs between forks are kept until one of the two interacting partners is lost due to merging of replication bubbles or replication of the chain ends. In this case, the Metropolis criterion is applied on *H*_*tot*_.

We investigate systematically the dynamical effects of introducing the non-trivial topology of the replication forks in the two scenarios explored in the main text (see Supplementary Note). In particular, we verify that the local dynamics of non-interacting sister forks (monomers with triple connectivity) and interacting sister forks (spring potential between two monomers with triple connectivity) agrees with theoretical predictions of local dynamics of the branching points in star polymers [77]. Furthermore, we include a comparison with a distinct implementation in Molecular dynamics for non-interacting sister forks. As usually observed in KMC and lattice simulations[72], the non-trivial topologies introduced by the forks lead to an artifactual decrease in the overall diffusion of the chain. We take great care of correcting for this by rescaling MCS time according to the current number of branching points in the system at a given time (See Supplementary Note).

#### 3. Single origin system

For every trajectory, a polymer of length *L* = 1001 is first initialized in a random unknotted configuration and evolved during 10^8^ MCS to reach a steady-state. Then, the single origin is fired and the replication proceeds as described above. Thanks to the previously described time mapping, we convert the most recent experimental estimation of replication fork speed of 2.2 kb/min [78] to *v* ≈ 10^−5^ monomer/MCS. In Fig. 2B, we also investigate *v* = 10^−4^ and 10^−3^ monomer/MCS.

For the static analyses of replication bubbles of different sizes, the origin is triggered and forks move very rapidly along the chain with *v* = 1 until the desired loop size is obtained. Then the system is evolved during 10^8^ MCS to reach steady-state.

For all the investigated conditions, we run 1000 simulated trajectories to collect enough statistics.

#### 4. Multiple origins system

The system is prepared as before but with *L* = 1000. In addition, now, a set of several origins is selected in order to obtain a regular, homogeneous distribution of firing positions along the chain. For all the investigated conditions, we run 1000 simulated trajectories to collect enough statistics (except for the 10 origins system where we averaged over 300 simulations) with *v* = 10^−5^ monomer/MCS (physiological speed).

The instantaneous replication case is obtained by using all the monomers as origins of replication and *v* = 1 for merging the resulting convergent replication forks.

#### 5. Simulations at different chain sizes

In Figure S12, we perform simulations for different chain sizes (*L* = 2000, 3000, 5000). For each case, we run 300 simulations to collect enough statistics. While increasing the polymer length, we kept the volumic fraction (Φ = 6%) and the inter-origin distance (i.e. one origin every 40 monomers) constant. For this analysis *J*_*sister*_ was set to 0 kT (no fork-fork interactions).

#### 6. Simulations with chain crossing

We introduce chain crossing moves in the lattice following the implementation by [68, 79]. Briefly, the system is initialized using the instantaneous replication mode (see above). After the chain being fully replicated, specific trial moves are introduced in the system allowing for chain crossing: a monomer *i* is randomly chosen and attempted to move to a random NN site; if the site is free, a standard trial move is attempted ; if the site is occupied (by a monomer *j*), the positions of *i* and *j* are exchanged if the new positions do not break chain connectivity. Such a move allows strand passing while conserving steric interactions. To collect enough statistics, we average over 300 simulations.

#### 7. Differential properties of replicated monomers

In Figure S13, we simulate a system where newly replicated monomers can self-interact. We opt for a copolymer approach as previously done for similar lattice models [21, 68, 72]. We include contact interactions between all the replicated monomers in the system (*status* = 1, − 1) with strength *Jpp*. The total energy of the system is thus characterized by an additional term *H*_*repl*_:

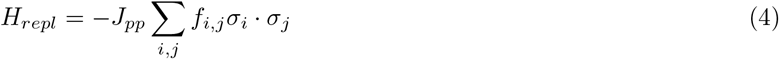

where *σ*_*i*_ = 1 if *status*(*i*) = 1, −1 otherwise *σ*_*i*_ = 0. *f*_*i*,*j*_ = 1 if monomers *i* and *j* are NN otherwise *f*_*i*,*j*_ = 0 .

In Figure S14, we simulate a system where newly replicated monomers have a different stiffness compared to the maternal chain (*l*_*k*_ = 100 nm). Concretely, when computing *H*_*bend*_ from Eq. 1), if monomer *i* is replicated (*status*(*i*) = 1, −1), *κ* is multiplied by a factor 0.5 (leading to *l*_*k*_ = 50 nm) or 1.38 (leading to *l*_*k*_ = 150 nm).

#### 8. Simulations at different volumic fractions

In Figure S15, we perform simulations for *L* = 1000 at different volumic fraction values Φ. We vary the box size in order to obtain Φ = 1%, Φ = 6% (standard concentration) and Φ = 25% while fixing *J*_*sister*_ = 100 kT (presence of fork-fork interactions). For each case, we run 300 simulations.

### B. Data analysis of simulations

#### 1. Computation of ⟨R_g_ ⟩ and ⟨k^2^⟩

To avoid any ambiguity do to the non-fixed number of monomer during replication we computed the average radius of gyration ⟨*R*_*g*_(*t*)⟩ and relative shape anisotropy ⟨*k*^2^(*t*)⟩ for an individual polymer copy. Therefore, prior to full replication we consider in the computation only the unreplicated monomers and those assigned to sister chromatid 1 (i.e with status = 1). For a given configuration, *R*_*g*_(*t*) and *k*^2^(*t*) are defined from the gyration tensor:

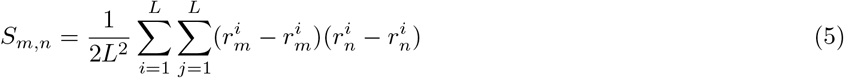

With 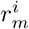 the *m* ∈ {*x, y, z*} coordinate of monomer *i*.

Given the principal moments of the gyration tensor *λ*_*x*_,*λ*_*y*_,*λ*_*z*_, the radius of gyration is defined as:

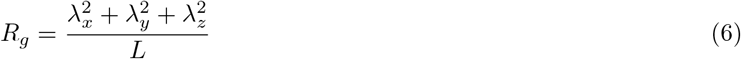

While the relative shape anisotropy:

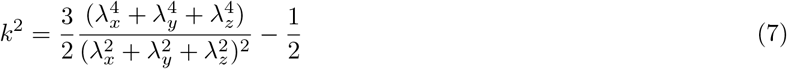

#### 2. Computation of contact maps

The contact probability between any two monomers *i* and *j* is defined as the probability that the 3D Euclidean distance *d*_*i*,*j*_ between *i* and *j* is less than a fixed radius of contact *r*_*c*_ = 50 nm.

For most of the computing contact maps during replication, we consider only intra-copy contacts, i.e. contacts between *i* and *j* belonging to the same chromatid. Note that before a monomer is replicated, we assume it to belong to both Sister chromatid 1 and Sister chromatid 2. Contacts between monomers with replication status +1 and −1 are not included. Pile-up plots in Fig. 4A and Fig. S4, S13 and S14 are computed by averaging the contact maps around the 25 origins in a time window of 2 ·10^5^ MCS, i.e. ≈ 40 sec.

We test if the patterns observed in pile-up plots are impacted by inter-chromatids contacts. In Figure S16, in a similar fashion to an HiC-like experiment, all the contacts between monomers are considered whatever their replication status. As a result, the observed contacts between *i* and *j* may originate from intra- and inter-copy contacts depending if *i* and *j* have been replicated at the time of computation of the map. Similar patterns are observed in this raw signal but with an enhanced background. Correcting such a time-dependent variation in monomer copy-number by dividing each element of the contact matrix *M* by the average copy-number *Cn* of *i* and *j* 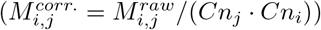 lead to similar pile-up plots than those obtained only with intra-copy contacts.

After full replication, we define as inter contacts only those occurring between monomers with different replication status, i.e. belonging to different sister chromatids. We obtain a symmetrical inter-contact matrix by combining the two different possibilities from a pair of positions:contacts between monomer *i* from chromatid 1 and monomer *j* from chromatid 2 and contacts between *i* from chromatid 2 and *j* from chromatid 1. Maps used in Fig. 6 and Fig. S8, S9, S10 and S12 at time *t* after replication, are defined by summing all contacts observed during a time window of 8 sec. The computation of average contact probability *P*_*intra*_(*s*) and *P*_*inter*_(*s*), plotting of matrices and pileup were done using the cooltools python library (v 0.5.1) [80, 81].

### C. Data analysis of experiments

#### 1. Data availability

The yeast Micro-C data used for Fig. 5 A from [13] was downloaded as cooler file from GEO GSE151553. The yeast Hi-C data used for Fig. 5B from [42] was downloaded as cooler file from GEO GSE162193. Data used in Fig. S7, was downloaded from https://makarich.fbb.msu.ru/agalicina/Lab_open/SCER/. All 3C data were balanced with ICE [82, 83] to correct for sequencing biases and copy number variation. The ChIP-Seq profile of Mcd1p was downloaded as bigwig file from GEO GSE151553. The microscopy data from Kitamura *et al*. used in Fig. 3B, was extracted directly from Figure 5A of [17].

#### 2. Pileup and averaged cohesin plots around origins and CARs in Saccharomyces cerevisiae

To define the early- and late-replicating origins, we used the “initiation probability landscape” signal (*IPLS*) from [40] (Fig.S5A). The *IPLS* is a signal which defines origin efficiency and is obtained through a machine learning approach in order to accurately reproduce experimental mean replication timing and Replication Forks Directionality data [78]. We use the Scipy function *find peak* [84] to extract the peaks of *IPLS* . We define the 15% highest peaks (signal > 5) as early origins of replication and all the remaining peaks whose signal is above 1 as late origins. Cohesin-associated regions (CARs) are defined by first binning at 1*kb* the Mcd1p ChIP-seq of Nocodazole arrested cells and then by using the Scipy function *find peak* [84] to extract the peaks. Early replicating CARs are CARs at a distance less than 40 kb from an early origins. Far from centromeres origins or CARs are defined by excluding those found at a distance lower than 40 kb from centromeres.

Pileup or average ChIP-seq plots in Fig. 5 and Fig. S5 were computed by aggregating the observed over expected experimental matrices using the coolpup python library [85] or by averaging the Mcd1p ChIP-seq profile at 1kbp resolution, respectively, around the set of loci of interest (origins or CARs).

### D. Code availability

Simulation codes is available at [86] under the branch name Repl2. The notebook associated with the analysis of 3C experiments is available at https://github.com/dariodasaro/HiC-Yeast-Data-analysis-around-origins.git

## ACKNOWLEDGMENTS

We are grateful to Aurèle Piazza, Benjamin Audit, Olivier Hyrien, Geneviève Fourel, Max Kolb, Elham Ghobadpour, Maria Barbi, Angelo Rosa, Davide Marenduzzo, Alexander Grosberg and the members of the Jost lab for fruitful discussions. We acknowledge Agence Nationale de la Recherche [DJ, MT: ANR-18-CE45-0022-01, ANR-21-CE13-0037-02; CV, DJ: ANR-21-CE45-0011-01] and ENS de Lyon (DD) for funding. We thank CBPsmn (Centre Blaise Pascal de simulation et modélisation numérique) of the ENS de Lyon for computing resources.

## Notes

### Competing Interest Statement

The authors have declared no competing interest.

### Summary of Updates

Our main revisions lie in (1) New simulations to investigate other parameters (like chromosome length, chromatin volumic density) or to test alternative scenarios (self-attraction, differential stiffness, strand crossing); (2) New analyses of experimental data to consolidate our conclusions; (3) An extensive rewriting of the text and supplementary material with new results, figures and discussions.

