## Supplementary Information for "DNA replication and polymer chain duplication reshape the genome in space and time"

[2] *École Normale Supérieure de Lyon, CNRS, Laboratoire de Physique, 46 Allée d'Italie, 69007 Lyon, France*

(Dated: June 17, 2024)

This PDF file includes:

- Supplementary figures S1 to S18
- Supplementary Notes
- URL links and Captions for Supplementary Videos S1 to S6

### I. SUPPLEMENTARY FIGURES

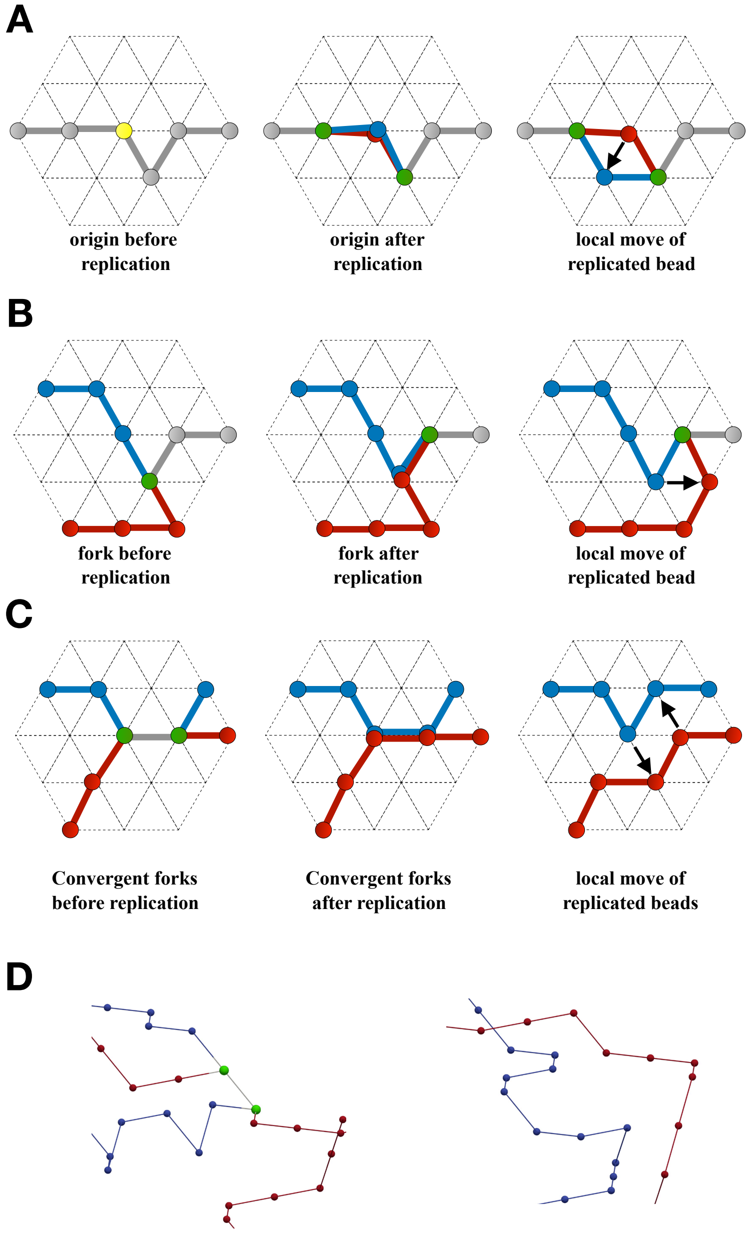


FIG. S1. **Scheme of replication moves in the lattice** (A) Monomer replication due to origin firing: (1) An origin (yellow) is in the unreplicated state. (2) A new monomer is introduced in the system and assigned to chromatid 2 (in red). Its position coincides with the site occupied by the old origin, now assigned to chromatid 1 (in blue), transiently violating the excluded volume rule. (3) The two monomer copies can then move in the lattice restoring the correct steric constraints. (B) Monomer replication due fork move: (1) Fork (green) prior to replication. (2) A new monomer is introduced in the system and assigned to chromatid 2. Its position coincides with the site occupied by the old fork, now assigned to chromatid 1. (3) The two monomer copies can move in the lattice restoring the correct steric constraints. (C) Steps following monomers replication due merging of bubbles: (1) Converging forks prior to replication. (2) Two monomers are introduced in the system and assigned to chromatid 2. Their positions coincide with the site occupied by the old forks, now assigned to chromatid 1. (3) The four monomer copies can move in the lattice restoring the correct steric constraints. (D) Snapshot of converging forks prior and following replication.

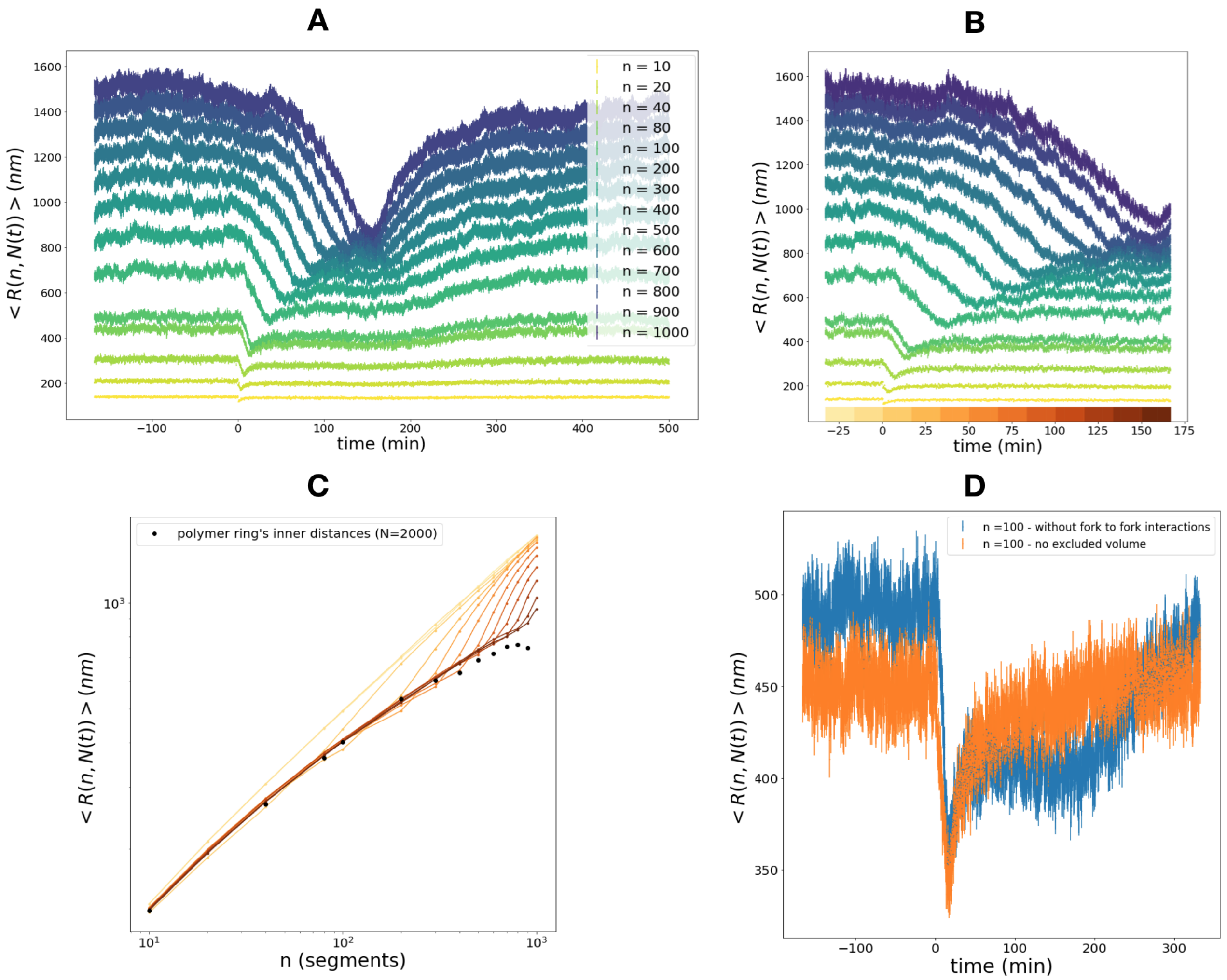


FIG. S2. **Spatial properties of a self-replicating polymer: single origin toy model without fork to fork interactions**. (A) 3D distance $\langle R(n, N(t))\rangle$ as a function of time (with $t = 0$ corresponding to the start of replication) for different pairs of monomers at a fixed distance $n$. (B) Zoom of panel (A). Colors below indicate discrete windows of time used to obtain the average $\langle R(n, N(t))\rangle$ at each $n$ in panel C. (C) $\langle R(n, N(t))\rangle$ in function of the linear distance $n$. At a given time interval $t^*$, a single curve is computed using $\langle R(n, N(t = t^*))\rangle$ for increasing values of n. Equivalently, each curve corresponds to a section of panel B on the y-axis. We observe a transition from linear-topology regime (lightest color) prior to replication to a more compact state just prior to bubble opening (darkest color). The black circles indicate the average $\langle R(n, N = 1000)\rangle$ within a polymer ring of size 2000 monomers at equilibrium. We show how the most compact state (darkest color, late replication) correlates well for short and intermediate scales ($n < 300$) with the internal distances of the equilibrated ring polymer. Lack of equilibration may be responsible for the minor compaction observed at larger scales (black points below the dark brown curve). (D) $\langle R(n, N(t))\rangle$ for $n = 100$ without sister-forks interactions in presence (blue) and absence (yellow) of excluded volume interactions. The result highlights how steric interactions are not necessary to observe the decrease in distance around the mean replication time of the two monomers. However, they are needed to maintain a significant compaction of their inner distances long after their replication.

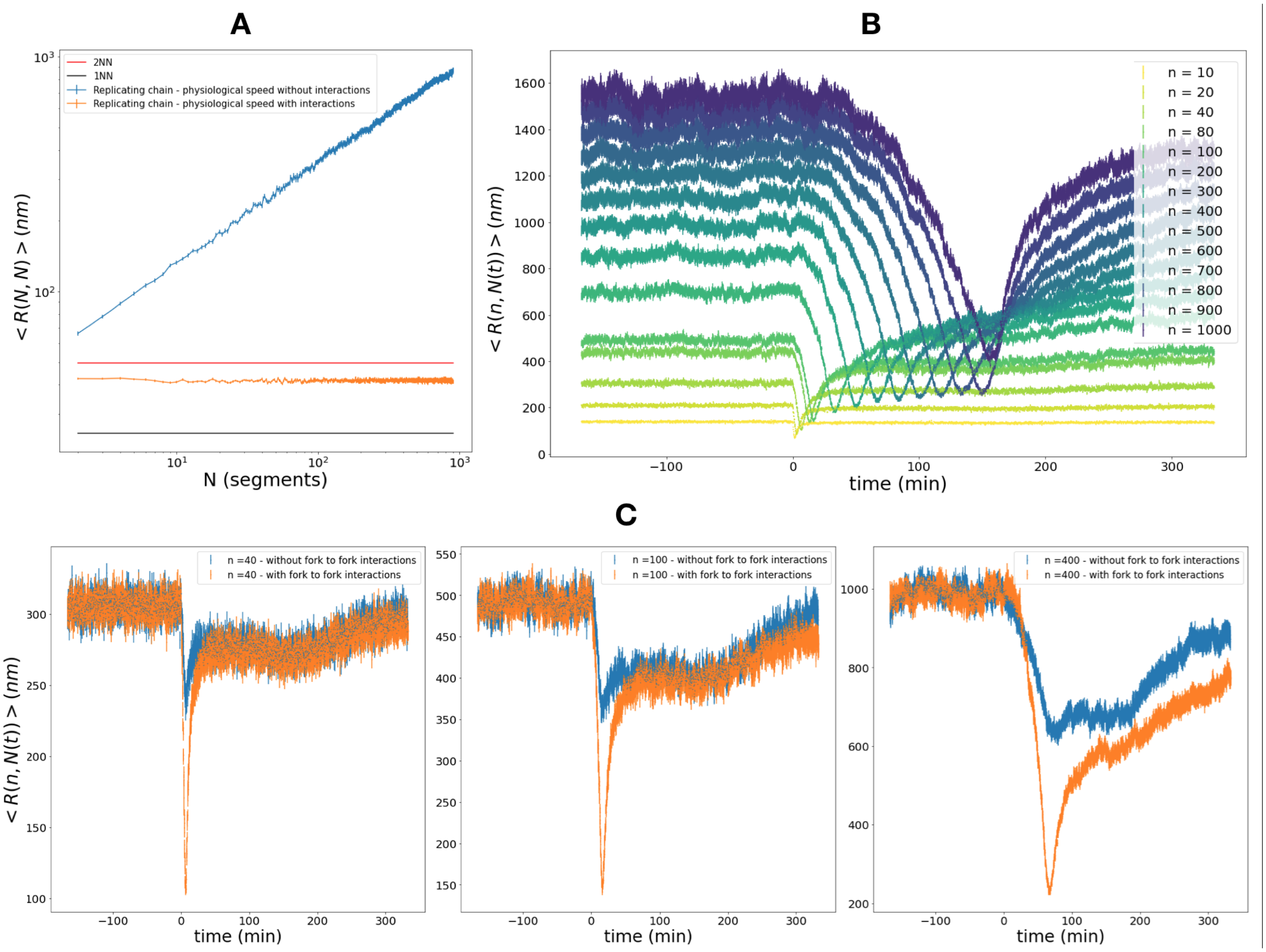


FIG. S3. **Spatial properties of a self-replicating polymer: single origin toy model with fork to fork interactions**. (A) Average fork-to-fork 3D distance $\langle R(N(t), N(t))\rangle$ as a function of the average number of replicated segments $\langle N(t)\rangle$ without (blue) and with (yellow) sister-forks interaction and at physiological speed. The black and red lines indicate the distance with the first and second nearest neighbours in the lattice (25 $nm$ and 50 $nm$). The spring potential used for sister-forks interaction (See Materials & Methods) saturates to a constant value (100 $kT$) when the forks are within a 3D distance of 50 $nm$. The $\langle R(N(t), N(t))\rangle$ for interacting forks (yellow curves) shows indeed how, for physiological replication speed, the interaction is maintained during their progression on the chain. (B) 3D distance $\langle R(n, N(t))\rangle$ as a function of time (with $t = 0$ corresponding to the start of replication) for different pairs of monomers at a fixed distance $n$ in presence of sister-forks interactions. (C) $\langle R(n, N(t))\rangle$ for $n = 40, 100, 400$ without (blue) and with (yellow) sister-forks interaction.

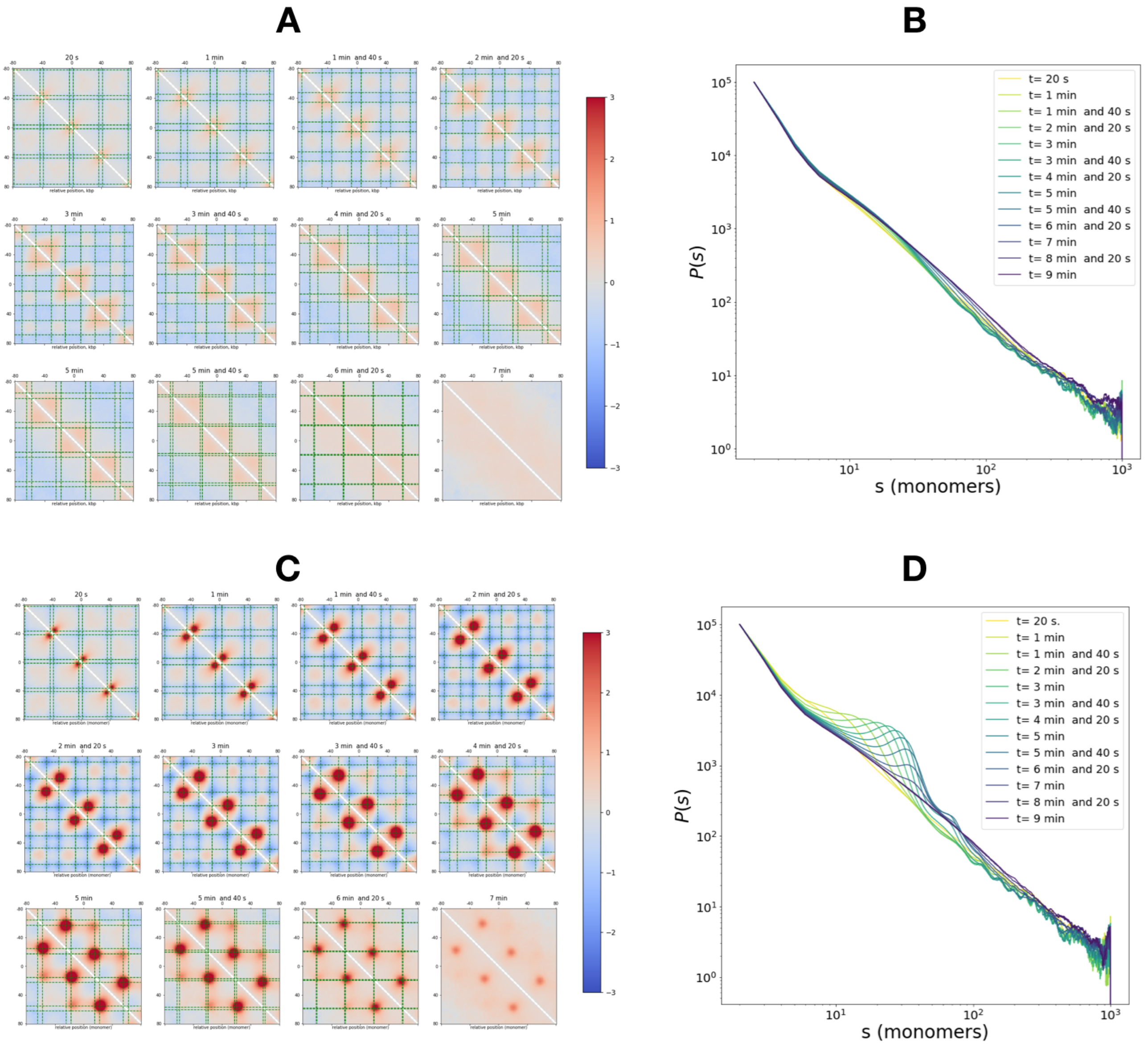


FIG. S4. **Features of contact maps for a 25 multi-origin system**. (A) Contact maps (for a radius of capture = 50 nm) at several time $t$ after replication started, averaged around an origin and normalized by the corresponding map in absence of replication. Dashed green lines indicate the average forks' position at this specific time step. (B) Average contact probability $P(s)$ between two monomers separated by a genomic distance $s$ for non interacting forks. The contact probability at the genomic distance $s$ corresponds to the average value of the s-diagonal of the contact map. (C) Same as A for interacting forks. (D) Same as B for interacting forks. The positioning and translocation in time of the contact loops in (C) are also clearly observable in the corresponding $P(s)$ curves.

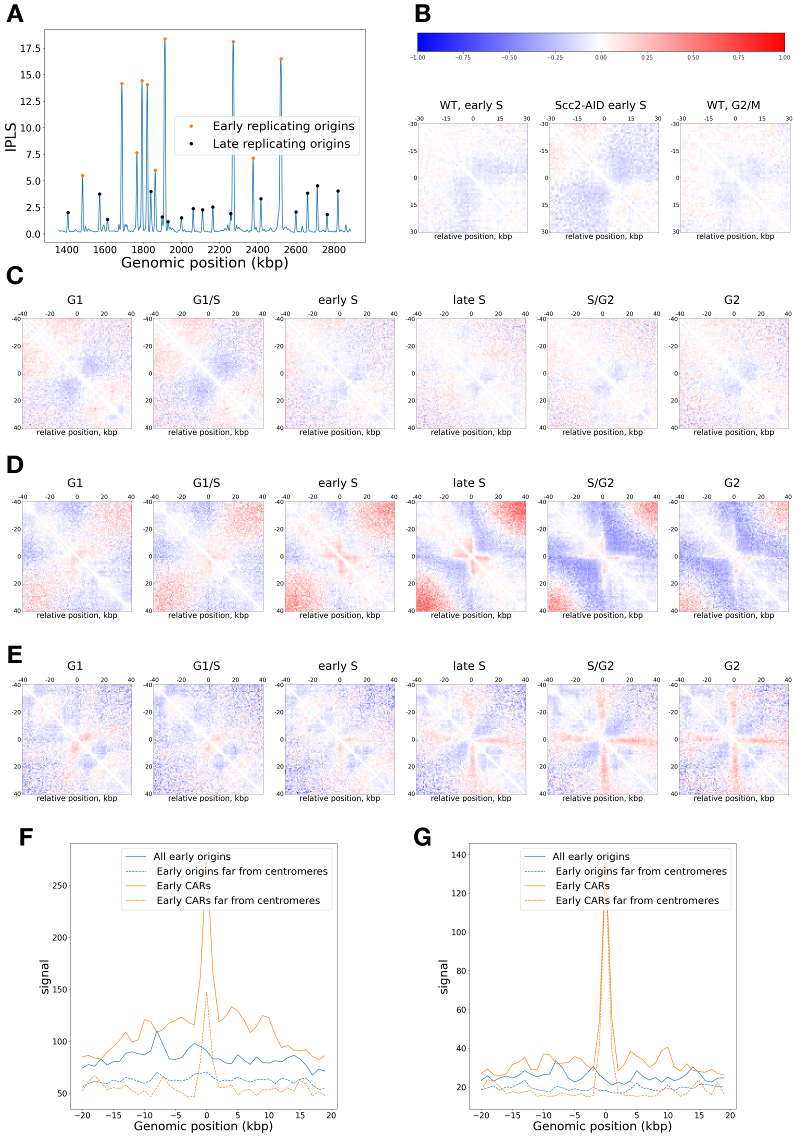


FIG. S5. **Experimental contact maps around origins and CARs in *Saccharomyces cerevisiae*.** (A) Example of the $IPLS$ signal for chromosome IV used to determine early and late origins of replication (See Materials & Methods). (B) Average normalized (Observed over Expected) contact maps at 1 kbp resolution around late replicating origins using experimental data from [1] resolution. The three plots show three different experimental conditions: Wild type (WT) cells arrested in Early S with HU treatment, scc2 depleted cells arrested in Early S with HU treatment and G2/M arrested cells. (C) Same as (B) but for experimental data from from [2] at different stages of the cell cycle and at a resolution of 800 bp. In both (B,C), no distinct contact pattern can be observed in all conditions.(D) Average normalized (Observed over Expected) contact maps at 800 kbp around early replicating CARs (See Materials & Methods). The contact pattern observed in G1 are reminiscent of the Rabl organization of yeast chromosomes during interphase as most of the early replicating CARS are close to centromeres. During S and G2, cross-like pattern appear, signature of CAR-CAR interactions mediated by cohesin [2]. (E) Same as (D) but excluding early replicating CARs within 40 kbp from centromeres. The Rabl-signature disappears but cross patterns are maintained. (F) Average ChIP-seq signal of cohesin subunit Mcd1p around origins and CARs in Early S. (G) Same as (F) but for Nocodazole-arrested cells (M-arrested). As expected, cohesin binds strongly at CARs but did not show any enrichment around early-replicating origins. For both (G) and (F) signals corresponding to CARs and early origins in chromosome XII were excluded from the computation.

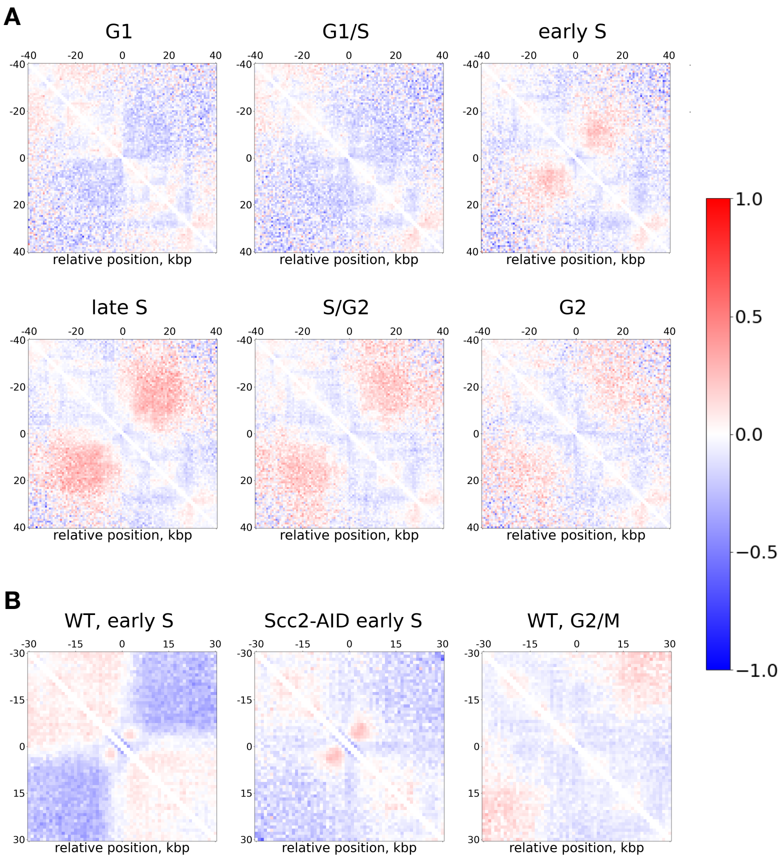


FIG. S6. **Experimental contact maps around strong origins of replication in *Saccharomyces cerevisiae* excluding sites within** 40 **kbp from centromeres**. (A) Average normalized (Observed over Expected) contact maps at 800 bp resolution around early replicating origins using experimental data from [2] at different stages of the cell cycle. (B) As in (A) but using experimental data from [1] at 1 kbp resolution. The three plots show three different experimental conditions: Wild type (WT) cells arrested in Early S with HU treatment, scc2 depleted cells arrested in Early S with HU treatment and G2/M arrested cells. For both (A) and (B), patterns are similar than those observed when all strong origins were included. They are even slightly more enhanced. The patterns (fountain-like) and their time evolution along cell-cycle are very consistent with the model predictions in the interacting-fork scenario.

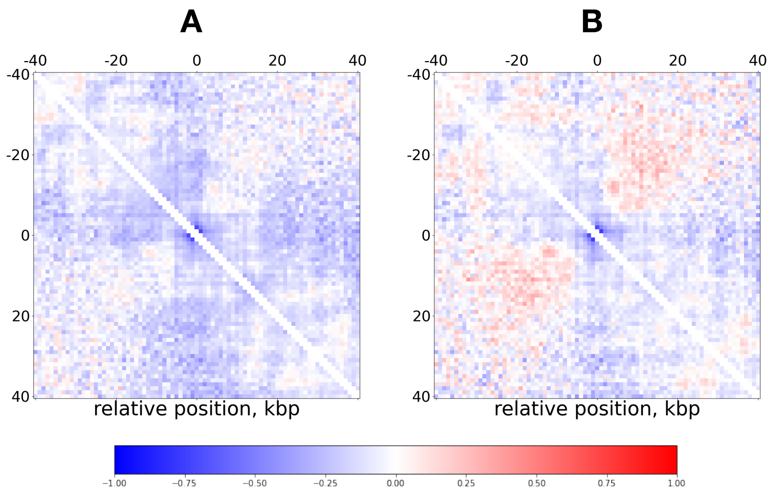


FIG. S7. **Experimental contact maps around strong origins of replication in *Saccharomyces cerevisiae*.** (A) Average normalized (Observed over Expected) contact maps at 1 kbp resolution around all early replicating origins using experimental data from [3] with cells arrested in S-phase. (B) As in (A) but excluding origins within 40 kbp from centromeres. The fountain-like pattern is only slightly visible when excluding origins far from the centromeres.

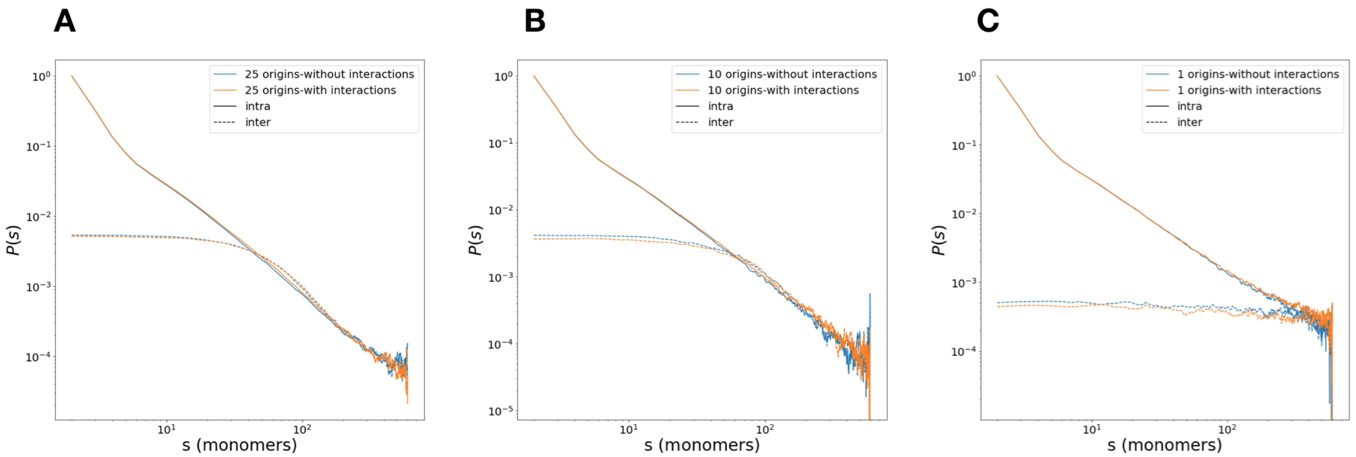


FIG. S8. **Effect of fork to fork interactions on intra vs inter** $P(s)$. (A) Average contact probability $P_{intra}(s)$ (solid lines) $P_{inter}(s)$ (inter) at $time = 0$ after replication for the 25 origin system, for non-interacting (blue) and interacting forks (yellow). (B) Same as (A) for 10 origins system. In both (A) and (B) the plots show how after all the merging events have occurred (full replication), there is no significant difference between the two interacting scenarios. This suggests that memory of the more compact and anisotropic looped structure is mostly lost and that interactions have little effect on the relative organization of the sister chromatids after replication. (C) Same as (A) for 1 origin system. The absence of a region where $P_{intra}(s) < P_{inter}(s)$ indicates that the two chains are not intertwined.

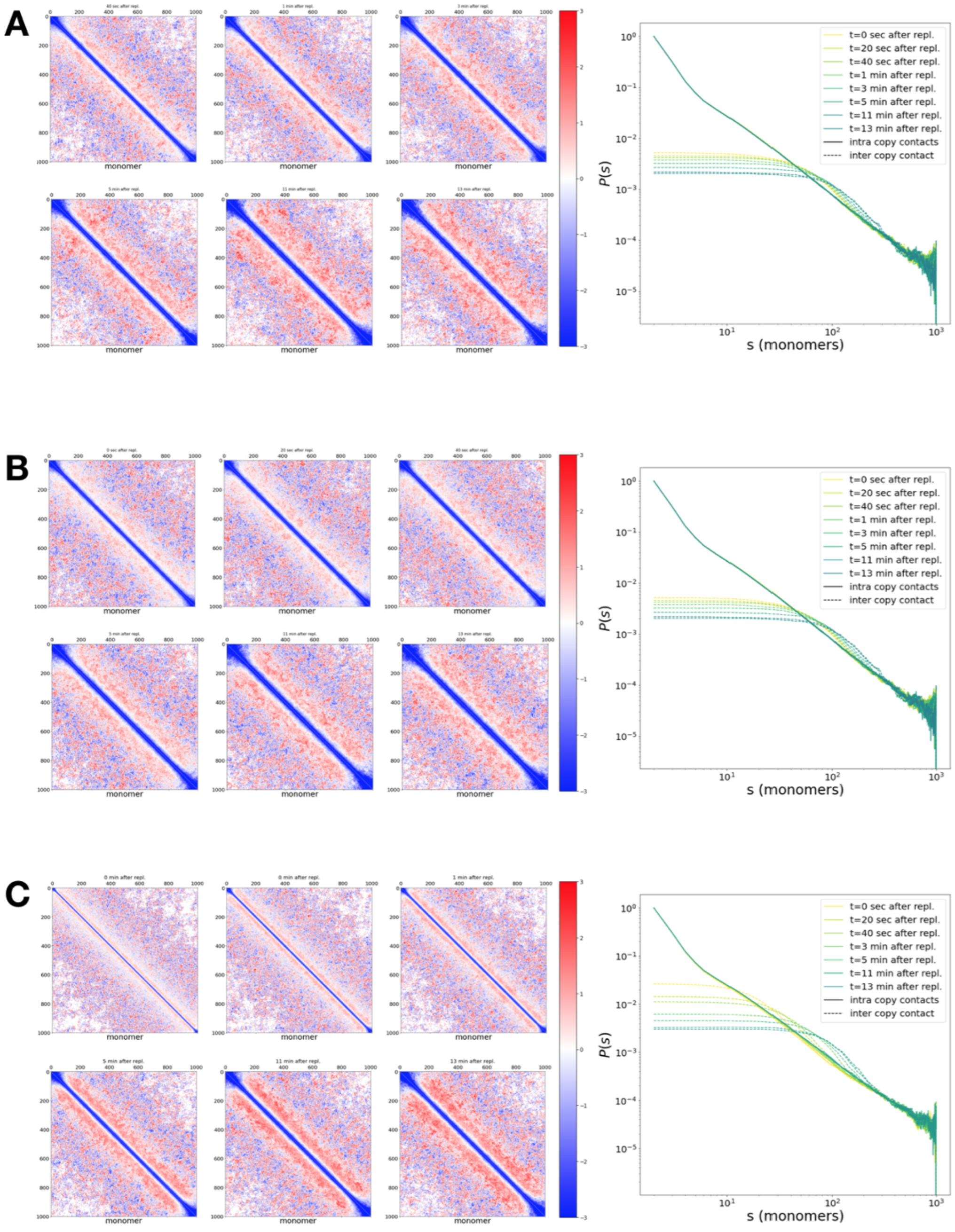


FIG. S9. **Relative organization of the sister chromatids: properties of a 25 multi-origin system**. (A) 3D organization of the 25 origin system without interaction. On the left: ratio between inter contact and intra contact maps at several times $t$ after replication. Intra maps only include the contacts between monomers of the same sister chromatid. Conversely, inter maps show the ones between monomers belonging to two different copies (See Materials & Methods). On the right: time series of intra (solid curves) and inter (dashed curves) contact probability $P(s)$ at a given time after full replication. As explained in more detail in the text, as the time progresses, the crossover value $s_{min}$ below which intra contact are higher (blue signal in the maps) and above which we find the region where $P_{intra}(s) < P_{inter}(s)$ (increasing red signal in the maps), gradually shifts towards greater values. The maps also highlight how in time, the two chains' end become progressively less entangled (triangular blue signal at the corner of the maps). (B) Same as A for 25 origins system with interaction. (C) Same as A,B for instantaneously replicating polymers.

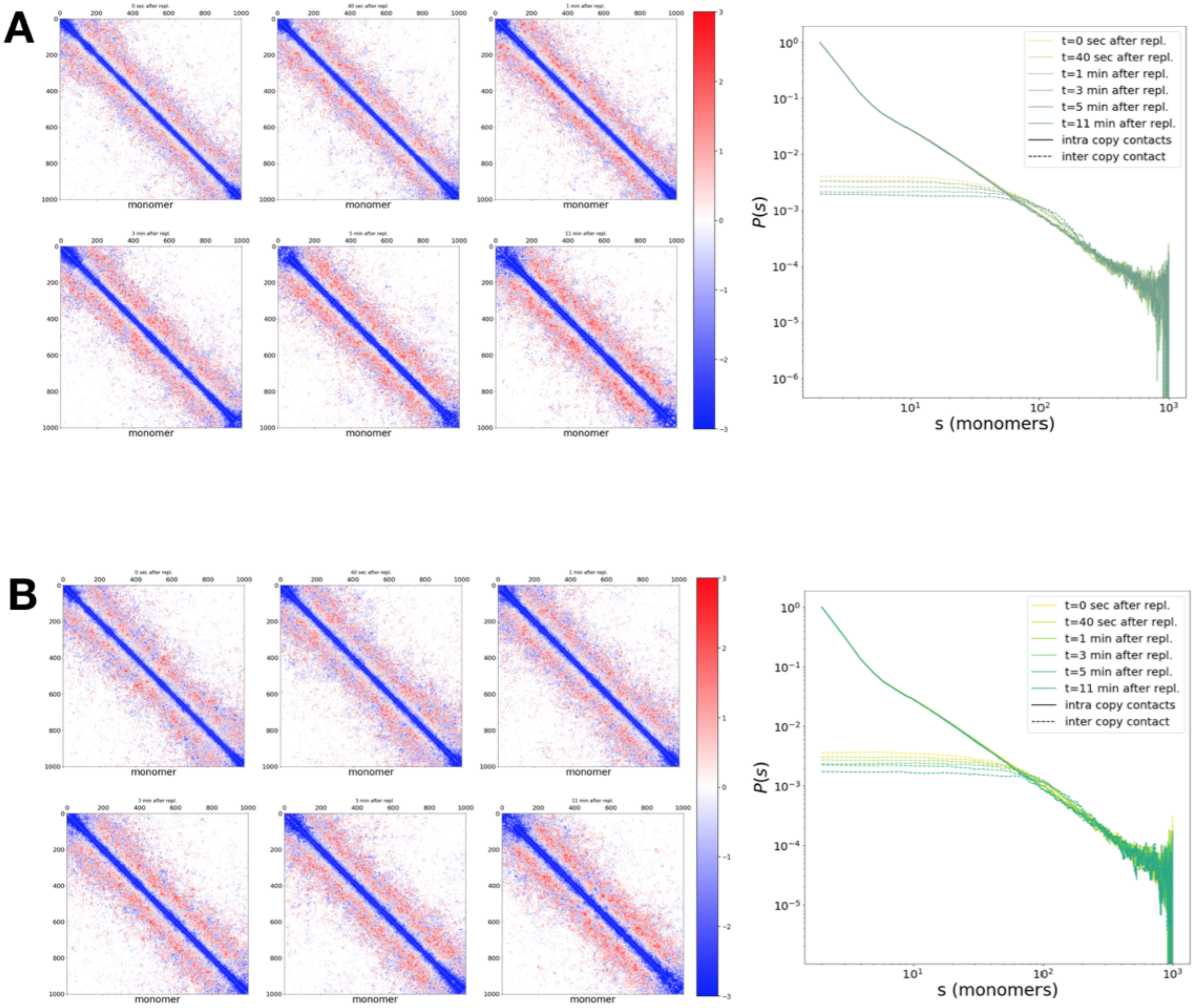


FIG. S10. **Relative organization of the sister chromatids: properties of 10 a multi-origins' system**. (A) 3D organization of the 10 origins system without interaction. On the left: ratio between inter contact and intra contact maps at several time steps $t$ after replication. Intra maps only include the contacts between monomers belonging to the same sister chromatid. Conversely, inter maps show the ones between monomers belonging to the two different copies (See Materials & Methods). On the right: time series of intra (solid curves) and inter (dashed curves) contact probability $P(s)$. As explained in more detail in the text, as the time progresses, the crossover value $s_{min}$ below which intra contact are higher (blue signal in the maps) and above which we find the region where $P_{intra}(s) < P_{inter}(s)$ (increasing red signal in the maps), gradually shifts towards greater values. The maps also highlight how in time, the two chains' end become progressively less entangled (triangular blue signal at the corner of the maps). (B) Same as A for 10 origins system with interaction.

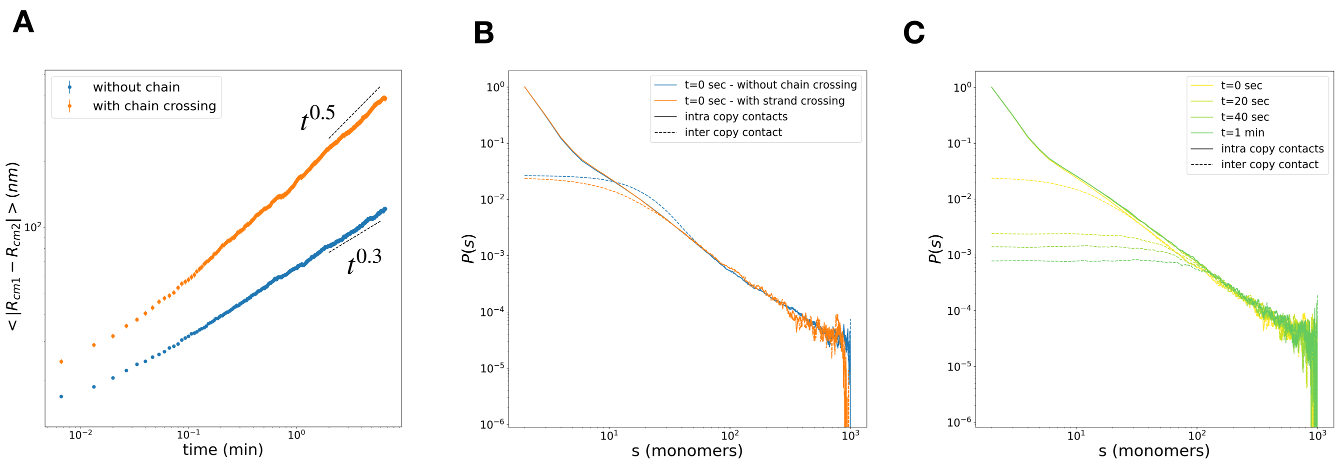


FIG. S11. **Simulations of instantaneous replication with and without chain crossing.** (A) Time evolution of the distance between the centers of mass of the sister chromatids $\langle |R_{cm2} - R_{cm1}|(t) \rangle$ in the limit of instantaneous replication in absence (blue) and presence (orange) of chain crossing. With strand passing, a diffusive regime is observed, faster than the subdiffusive regime observed in presence of topological constraints. (B) Intra (solid curves) and inter (dashed curves) contact probability $P(s)$ at time=0 after replication (first 8 seconds) in absence (blue) and presence (orange) of chain crossing. (C) As in (B) but at different times after the end of replication for the chain crossing simulation. The results in (B) and (C) suggest how, in presence of chain crossing, we loose the shoulder in the region of $P_{intra}(s) < P_{inter}(s)$ and that the relaxation is much faster in presence of chain crossing. These findings are consistent with the proposed model where, in absence of chain crossing, the topological constrains arising from the intertwined structure can only be relieved by relative motion of the two chains.

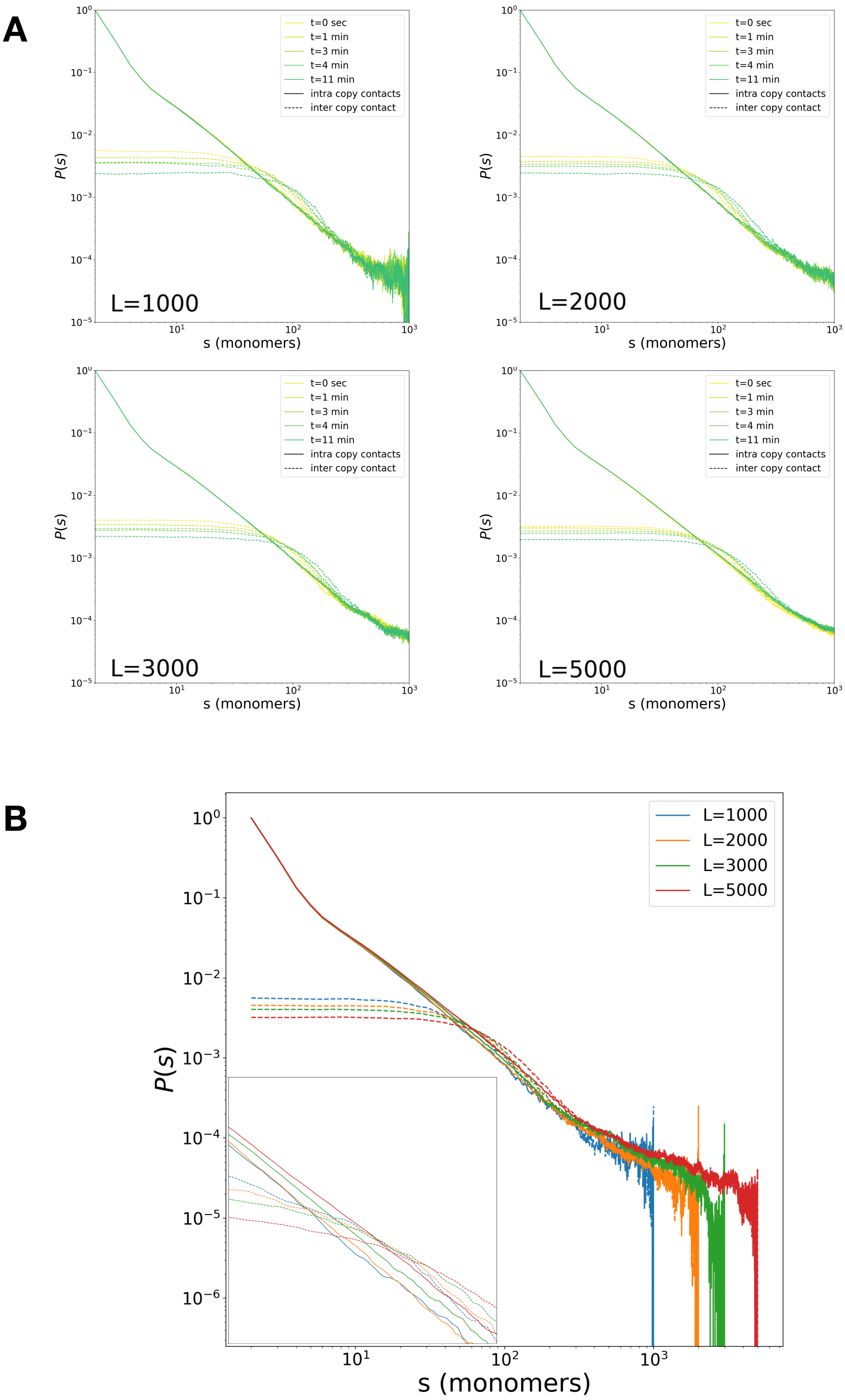


FIG. S12. **Relative organization of the sister chromatids for different chain sizes $L$ at a constant origin density**. (A) Intra (solid curves) and inter (dashed curves) contact probability $P(s)$ at different $t$ after full replication for different values of the chain size $L$. (B) Intra (solid curves) and inter (dashed curves) contact probability $P(s)$ at $t = 0$ after full replication for different $L$. (Inset) Zoom over the region where $P_{intra}(s)$ and $P_{intra}(s)$ intercept ($s_{min}$). The result show systematically bigger $s_{min}$ for increasing $L$ and systematically lower contacts for $s < s_{min}$.

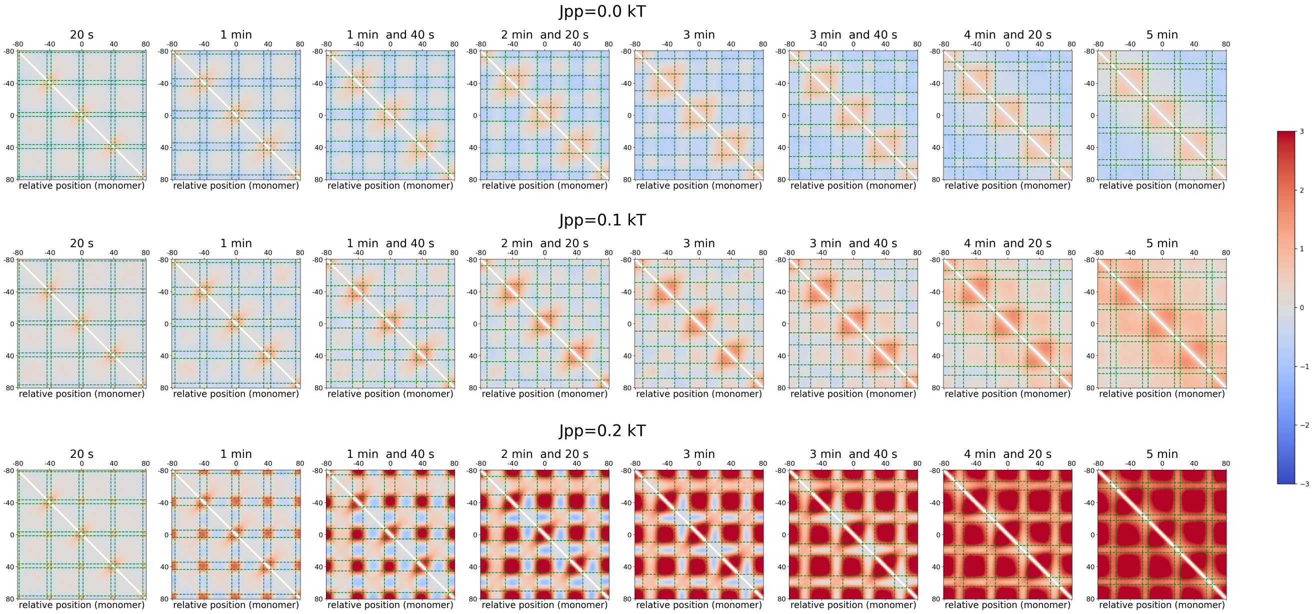


FIG. S13. **Features of contact maps for a 25 multi-origin system with self-attraction between newly replicated monomers**. Contact maps (radius of capture = 50 nm) at several time $t$ after replication started, averaged around an origin and normalized by the corresponding map in absence of replication. The analysis was done for three different strengths $Jpp$ of the self-attraction between newly replicated monomers in absence of fork-fork interactions: $Jpp = 0.0$ (top), 0.1 (middle), 0.2 kT (bottom). As bubble size increase in time and $Jpp$ is augmented, the cooperative effect of the interacting monomer result in stronger interactions within and between replication bubbles. The arising pattern at the bubble level (green square around origins) resemble the one of non-interacting forks rather than the looped pattern described in the text for interacting forks.

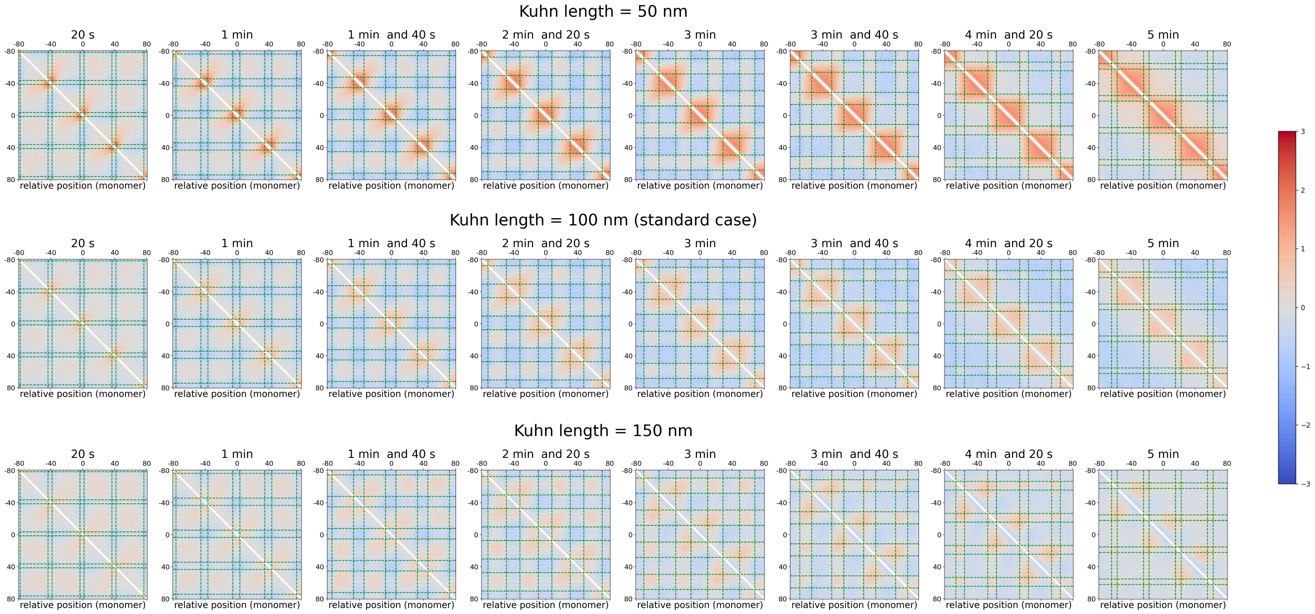


FIG. S14. **Features of contact maps for a 25 multi-origin polymer for different rigidities of the replication bubble**. Contact maps (radius of capture = 50 nm) at several time $t$ after replication started, averaged around an origin and normalized by the corresponding map in absence of replication. The analysis was repeated for three Kuhn length $l_K$ for the newly replicated monomers and in absence of fork-fork interactions: $l_k$ = 50 (top), 100 (middle) and 150 nm (bottom). The middle case correspond to the standard model where the stiffness of replicated and unreplicated regions of the polymer is the same. As expected, simulations show increase of contacts for softer polymers and depletion for stiffer one. Overall, the results do not exhibit fountain-like patterns.

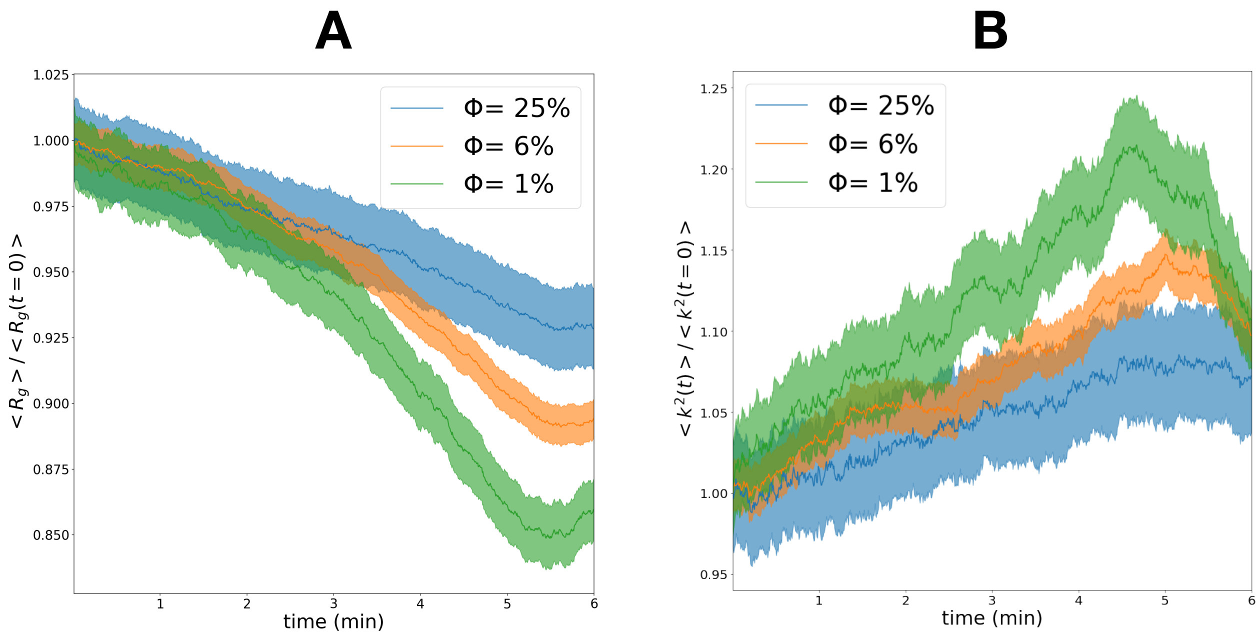


FIG. S15. **Large-scale organization is impacted by the volumic fraction in the interacting fork scenario.** (A) Time-evolution of the average radius of gyration $\langle R_g(t)\rangle$ normalized by $\langle R_g(t=0)\rangle$ at different volumic fractions $\Phi$. (B) Time-evolution of the relative shape anisotropy $\langle k^2(t)\rangle$ normalized by $\langle k^2(t)\rangle$ at different volumic fractions $\Phi$. Qualitatively, the same effects are observed whatever the density: the radius of gyration exhibits an optimal decrease and the anisotropy is maximal just before the end of replication. But quantitatively, their strengths are larger for lower densities.

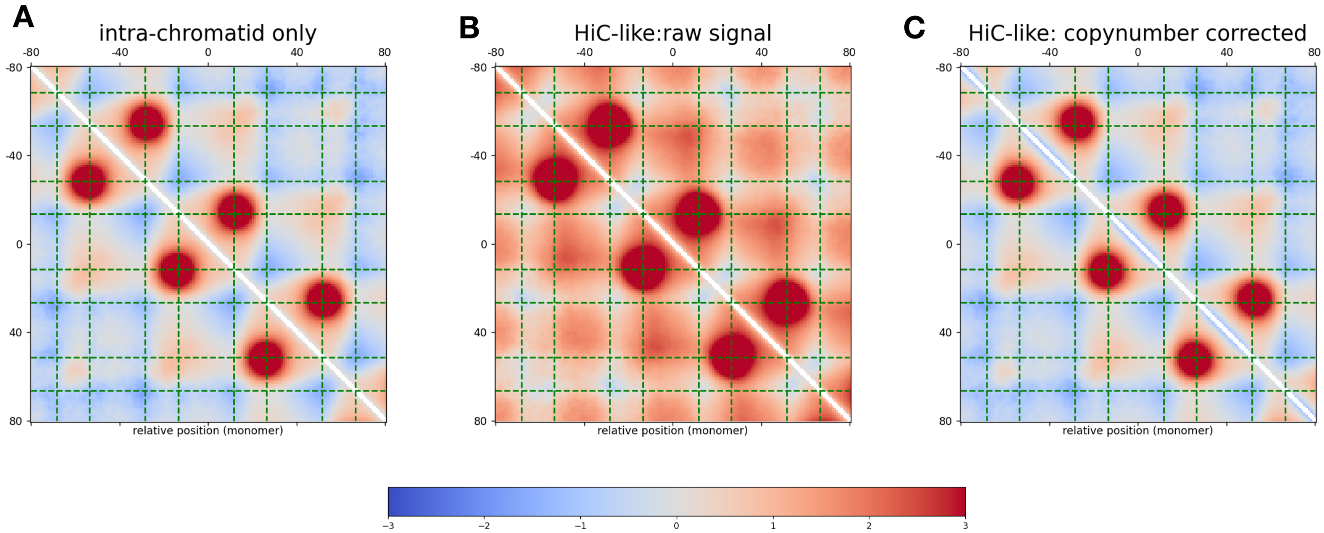


FIG. S16. **Computation of contact maps during replication** (A,B,C) Average contact maps around origins at time=3 min and 40 seconds predicted by the model in the interacting fork scenario.(A) Computation using only intra-chromatid contacts as in the main text. (B) Computation using both intra and inter total number of contacts (See Materials & Methods). We can observe stronger contacts between distal replication bubbles which are due higher average copy numbers around origins leading to a greater number of contacts assigned to the replicated regions. (C) Same as (B) but normalized by accounting for the average copy-number of each genomic position at the given time (See Materials & Methods). Such normalization recover the main structural feature observed in (A). This condition should mimic the experimental data which integrate both intra- and inter-chromatid contacts and are balanced using ICE, a method that effectively normalized for copy-number.

### II. SUPPLEMENTARY NOTES

#### 1. Dynamics of the forks

Triple connectivity at the forks constitutes a strong local topological constraint. Following our standard on-lattice kinetic Monte-Carlo (KMC) algorithm [4], to move a fork, a random trial position is picked among the nearest neighbor lattice sites and is rejected if it breaks the triple connectivity at the fork and if the site is already occupied. Equivalently to the standard moves in the linear part of the chain, this new specific move does not break detailed balance. To test if introducing these topological constraints in our KMC framework may affect the chain dynamics, we compared our implementation to theoretical predictions described in [5] and simulations using Molecular Dynamics (MD) (see below).

First, we study the dynamics of a single fork by implementing a simple star polymer constituted by three branches of equal size with 100 segments per branch (nobending rigidity and volumic fraction= 1%). We computed the Mean Square Displacement (MSD) $g_1$ ($\equiv \langle \overrightarrow{r_\alpha}(t+\Delta t) - \overrightarrow{r_\alpha}(t) \rangle_t$) for each monomer located at a given linear distance $\alpha$ from the branching point and compared the result to an equivalent implementation in MD simulations (Fig. S17 A). For comparison between KMC and MD, we rescale KMC and MD times to impose $g_1^{KMC}(\alpha = 50, t) \approx g_1^{MD}(\alpha = 50, t)$.

At small timescales, we find very good agreement between KMC and MD with $g_1 \sim t^{0.5}$ and a decrease in mobility as we approach from the branching point. This suggest that at short time scale, our KMC does not lead to strong dynamical artifacts. To confirm that, we simulate a larger system (three branches of equal size, each consisting of 500 segments)(volumic fraction=5%) and found that at a small timescales, the decrease in mobility at the branching point observed in KMC is also consistent with the theoretical model [5] (Fig. S17 B).

However at larger timescale, in Fig.S17 A, we observed a significant deviation between KMC and MD for small $\alpha$. This suggest a possible problem in the global diffusion of the polymer chain. Indeed for longer time-scale $g_1$ converges to the MSD of the center of mass of the polymer, $g_3(t) = \langle \overrightarrow{r}_{\rm cm}(t+\Delta t) - \overrightarrow{r}_{\rm cm}(t) \rangle_t$. In fact, since we are simulating an isolated system, $g_3$ should only depend on the total number of monomers [6] and be independent of the presence of branching points within the polymer topology. The observed decrease of $g_3$ in the presence of branching points is actually a standard artifact in KMC and lattice algorithms [7]. This can be corrected by "rescaling" time according to the number of branching points in the system at a given time, ie the time mapping between KMC and real time is now dependent on the current topology of the chain.

To estimate this correction, we consider a simple system with one replication loop of total size 100 monomers and two dangling ends of 50 monomers each (norigidity, volume fraction=0.9%). We compute $g_3$ and estimate the number of extra trial moves $corr_{free\_fork}$ per free branching point to rescue the expected $g_3$ of a linear polymer of the same size (200 monomers)(Fig. S17 C): $MCS = N_{monomer} + \boldsymbol{corr_{free\_fork}} \cdot N_{free\_fork}$ with $N_{monomer}$ the number of monomers and standard number of trial moves per Monte-Carlo step (MCS) and $N_{free\_fork}$ the number of free forks in the considered topology. We find an optimal value of $corr_{free\_fork} = 6$ for free — non interacting — forks. We demonstrate that this correction is additive by using the same correction in several topologies with 700 monomers (volumic fraction=3%) with an increasing number of forks by consistently recovering the expected behavior for $g_3$ (Fig. S17 D).

We expect that interaction between sister forks will result in even greater perturbation of in fork mobility due to stronger topological constraints. Computing $g_1$ at the branching point for the same toy topology with 200 monomers with a single-bubble with $Jsister = 100$, $threshold = 25$nm and volumic fraction=0.9% (see Materials and Methods), we recover a decrease in mobility qualitatively comparable to the prediction of a branching point of a star polymer with 6 arms (two connected triple connectivity monomers) (Fig. S18 A). In the main text, we use a $threshold = 50$nm which qualitatively retains comparable local dynamics while limiting the decrease of in global mobility (Fig. S18 B). However, even for $threshold = 50$ nm, artifacts on the global diffusion of the polymer can be observed. Computing $g_3$ for the same toy model, we find that for interacting sister forks, an optimal value of additional Monte Carlo moves $corr_{interac\_fork} = 17$ rescues the expected behavior (Fig. S18 C). The correction appears also to be additional and thus appropriate to have a runtime correction according to the current number of sister forks in the system at a given time (Fig. S18 D).

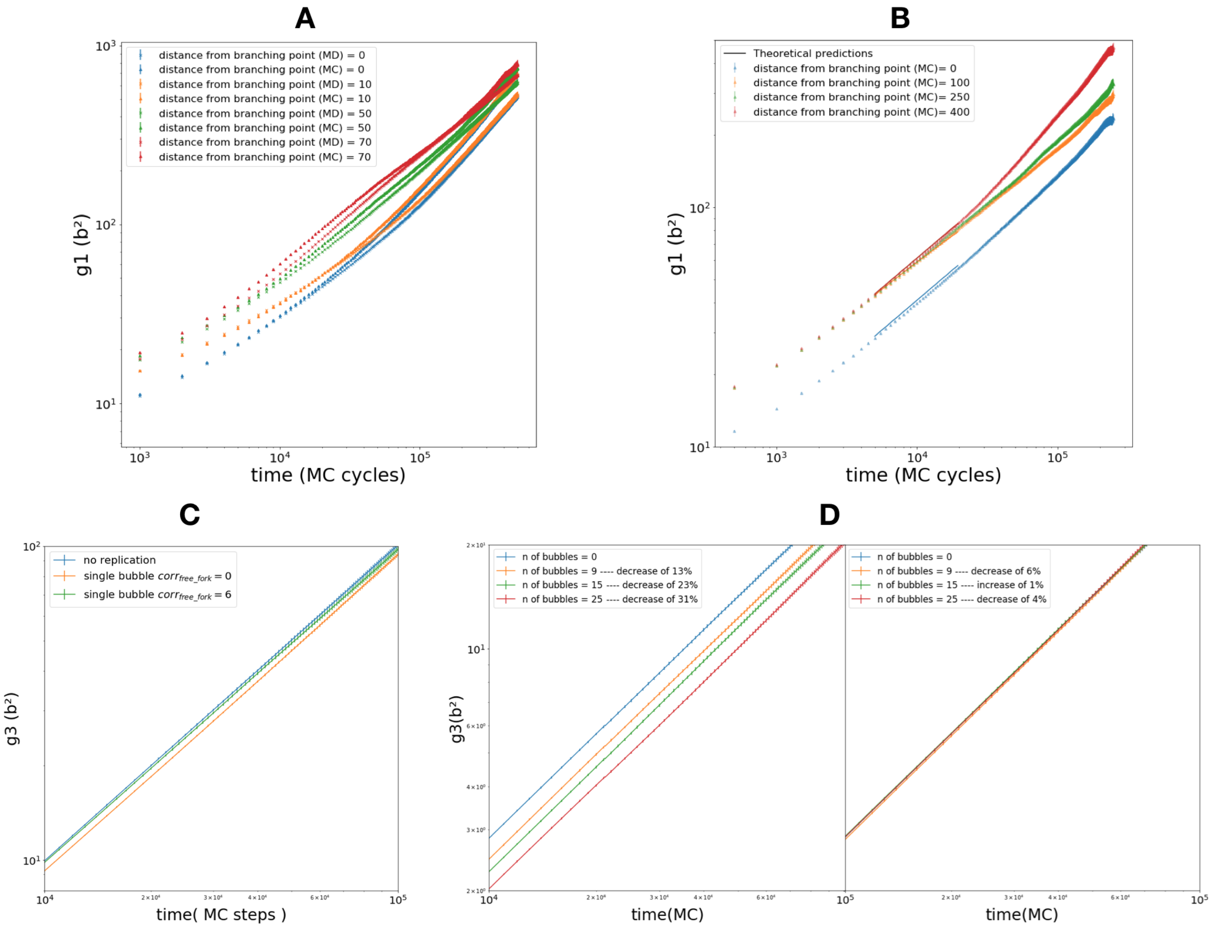


FIG. S17. **Reduced mobility at free branching points** (A) $g_1$ as a function of the distance $\alpha$ from the branching point for a 301 monomers star polymer computed in the lattice and with molecular dynamics simulations. (B) $g_1$ as a function of the distance from the branching point for a 1501 monomers star polymer computed in the lattice. Solid lines correspond to the theoretical expected decreases in function of distance from the branching point. (C) Decrease in $g3$ in presence of replication (blue curve and yellow curve) forks for a single-bubble symmetrical system of 200 monomers. Rescue of the expected behavior by $corr_{free\_fork} = 6$ moves for each fork (green curve). (D) Left panel: $g_3$ for several 700 beads systems with an increasing number of replication bubbles. In blue the corresponding $g_3$ for a linear chain containing the same number of monomers (blue curve). Right panel: same as left panel in presence of extra corrective moves.

Overall, for any current topology present in our simulations, the number of trial moves is corrected as:

$$MCS = N_{monomer} + \boldsymbol{corr_{single\_fork}} \cdot N_{single\_fork} + \boldsymbol{corr_{interac\_fork}} \cdot N_{interac\_fork} \tag{1}$$

#### 2. *Details of MD simulations*

MD simulations of isolated forks were run by considering a symmetric star polymer comprised of 3 identical branches, each made up of 100 monomers, and joined at their extremities. Calculations were run in periodic boundary conditions in a cubic box, whose linear dimension was chosen such that the polymer packing fraction is of order $\sim 0.001\,\%$ so as to neglect the effects of molecular crowding. Chain connectivity was imposed through a standard FENE spring potential, while local excluded volume was implemented via a truncated and shifted Lennard-Jones pairwise contribution (see

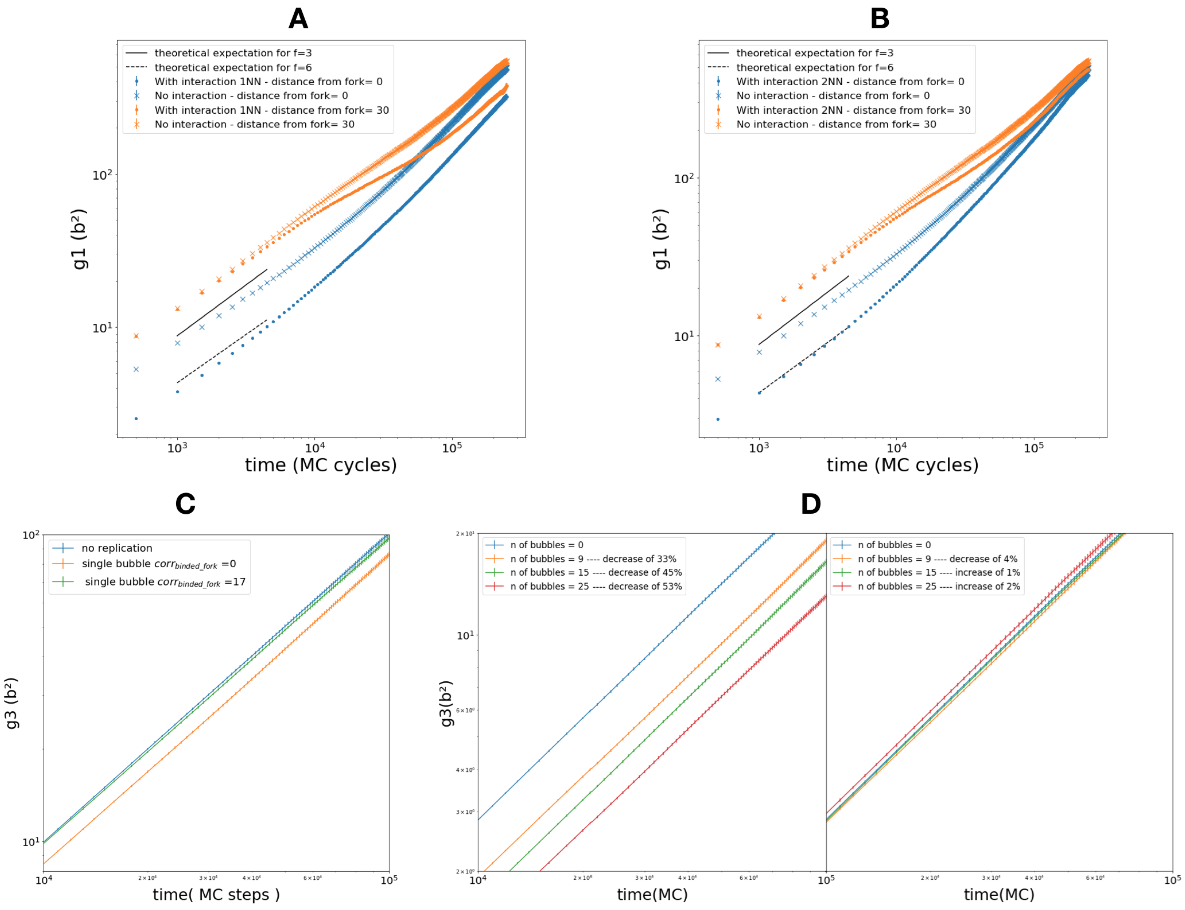


FIG. S18. **Reduced mobility for interacting branching points** (A) $g_1$ in function of the distance $\alpha$ from the branching point for a 200 monomers single-bubble system. The blue curve shows $g_1(\alpha)$ in absence of fork to fork interaction while the yellow curve in presence of interactions with a $threshold = 25$nm. Solid lines correspond to the expected theoretical decreases at the branching point with $f = 3$ and $f = 6$ arms. (B) Same as A but for $threshold = 50$nm. (C) Decrease in $g_3$ in presence of free replication (blue curve and yellow curve) forks for a single-bubble symmetrical system. Rescue of the expected behavior by $corr_{binded\_fork} = 17$ moves for each fork (green curve). (D) Left panel: $g_3$ for several 700 beads systems with an increasing number of interacting replication bubbles. In blue the corresponding $g_3$ for a linear chain of equal size (blue curve). Right panel: same as left panel in presence of extra corrective moves.

[8] for the full numerical details of the parameters used). Simulations were conducted in the canonical ensemble using a Langevin thermostat, and were performed based on the HOOMD-blue molecular simulation package [9]. Conformations were initially relaxed over the course of $\sim 10^7$ integration time steps, following which production runs of $\sim 10^8$ MD steps were used for data collection.

### III. SUPPLEMENTARY VIDEOS

- **Video S1**: `https://youtu.be/7gcjG_wRRbU`
  Scheme of the self replicating polymer. On the right the various steps of the 1D replication dynamics while on the left its 3D realization inside the lattice.

- **Video S2**: `https://youtu.be/mGJOefvrQuI`
  Example of two non-interacting and interacting sister forks following the firing of a origin.

- **Video S3**: `https://youtu.be/9OkoOF7E_zA`
  "Yeast-like" chromosome replicating from 25 origins regularly inter-spaced along the chain with sister forks' interactions.

- **Video S4**: `https://youtu.be/7ISEUoPMHjg`
  Contact maps (radius of capture = 50 nm) at several timestep $t$ after beginning of replication, averaged around an origin and normalized by the corresponding map in absence of replication. Dashed green lines indicate the average forks' position at this specific time step.

- **Video S5**: `https://youtu.be/31UauSfL66w`
  Intertwined sister chromatids with loose ends at $t = 13$ min after the full replication in the case of non-interacting forks.

- **Video S6**: `https://youtu.be/4EWXwepuL1Q`
  Ratio between inter contact and intra contact maps at several time steps $t$ after replication . Intra maps only include the contacts between monomers belonging to the same sister chromatid. Conversely, inter maps show the ones between monomers belonging to the two different copies (See Materials & Methods).
